# Comparing newly developed SNP barcode panels with microsatellites to explore population genetics of malaria parasites in the Peruvian Amazon

**DOI:** 10.1101/2024.09.09.611954

**Authors:** Luis Cabrera-Sosa, Mahdi Safarpour, Johanna H. Kattenberg, Roberson Ramirez, Joseph Vinetz, Anna Rosanas-Urgell, Dionicia Gamboa, Christopher Delgado-Ratto

## Abstract

Malaria molecular surveillance (MMS) can provide insights into transmission dynamics, guiding national control/elimination programs. Considering the genetic differences among parasites from different areas in the Peruvian Amazon, we previously designed SNP barcode panels for *Plasmodium vivax* (Pv) and *P. falciparum* (Pf), integrated into AmpliSeq assays, to provide population genetics estimates of malaria parasites. These AmpliSeq assays are ideal for MMS: multiplexing different traits of interest, applicable to many use cases, and high throughput for large numbers of samples. The present study compares the genetic resolution of the SNP barcode panels in the AmpliSeq assays with widely used microsatellite (MS) panels to investigate Amazonian malaria parasites.

Malaria samples collected in remote areas of the Peruvian Amazon (51 Pv & 80 Pf samples) were characterized using the Ampliseq assays and MS. Population genetics estimates (complexity of infection, genetic diversity and differentiation, and population structure) were compared using the SNP barcodes (Pv: 40 SNPs & Pf: 28 SNPs) and MS panels (Pv: 16 MS & Pf: 7 MS).

The genetic diversity of Pv (expected heterozygosity, *He*) was similar across the subpopulations for both makers: *He*_MS_ = 0.68 - 0.78 (p = 0.23) and *He*_SNP_ = 0.36 - 0.38 (p = 0.80). Pairwise genetic differentiation (fixation index, F_ST_) was also comparable: F_ST-MS_ = 0.04 - 0.14 and F_ST-SNP_ = 0.03 - 0.12 (p = 0.34 – 0.85). No geographic clustering was observed with any panel. In addition, Pf genetic diversity trends (*He*_MS_ = 0 – 0.48 p = 0.03 – 1; *He*_SNP_ = 0 - 0.09, p = 0.03 – 1) and pairwise F_ST_ comparisons (F_ST-MS_= 0.14 – 0.65, F_ST-SNP_ = 0.19 – 0.61, p = 0.24 – 0.83) were concordant between the panels. Similar population structure clustering was observed with both SNP and MS, highlighting one Pf subpopulation in an indigenous community.

The SNP barcodes in the Pv AmpliSeq v2 Peru and Pf AmpliSeq v1 Peru assays offer comparable results to MS panels when investigating population genetics in Pv and Pv populations. Therefore, the AmpliSeq assays can efficiently characterize malaria transmission dynamics and population structure and support malaria elimination efforts in Peru.

## 1 Introduction

In 2022, nearly 250 million malaria cases were estimated worldwide, mainly concentrated in low and middle-income countries of Africa, Latin America (LATAM) and Southeast Asia (World Health Organization, 2023). Many countries are making efforts to eliminate malaria in the next years by implementing National Malaria Control and Elimination Programs (NMCP/NMEPs).

The World Health Organization (WHO) states in the Global Strategy 2016 – 2030 that transforming malaria surveillance into a key intervention in NMCP/NMEPs is a crucial pillar for malaria elimination (World Health Organization, 2021). Surveillance can strengthen NMCP/NMEPs by providing information on transmission dynamics, drug resistance evolution, etc., that can be used to propose strategies for control and elimination and inform decision-making (Dalmat et al., 2019).

Malaria molecular surveillance (MMS) can support traditional epidemiologic surveillance based on microscopy or rapid diagnostic tests (RDT) to understand transmission dynamics (Golumbeanu et al., 2023). MMS also has the potential to be implemented into NMCP/NMEPs to contribute to the disease control efforts (Noviyanti et al., 2020). Scenarios where genetic surveillance data of *Plasmodium vivax* (Pv) and *P. falciparum* (Pf) parasites can inform malaria strategies have been identified (Dalmat et al., 2019). In that sense, population genetics can be used for these “use cases” to measure transmission intensity (Pava et al., 2020), identify parasite relatedness and gene flow (Brown et al., 2021; Schaffner et al., 2023), distinguish imported cases and transmission chains (Trimarsanto et al., 2022), characterize outbreaks and transmission foci (Wasakul et al., 2023) and determine genetic diversity and population structure (Nderu et al., 2019; Sy et al., 2022).

Microsatellites (MS) panels have been used extensively to study *Plasmodium* population genetics. MS are high polymorphic, short tandem repeat loci found in approximately 10% of the *Plasmodium* genome (Mathema et al., 2020). The neutral MS panels (not under selection pressure) are ideal for determining the population’s genetic changes (Anderson et al., 2000; Ferreira et al., 2007; Imwong et al., 2007).

More recently, single nucleotide polymorphisms (SNPs) have gained popularity for population genetics analysis, not only for malaria but also for other infectious diseases and in conservation studies (Brashear and Cui, 2022; Theissinger et al., 2023). Although SNPs have low allelic diversity, they have high prevalence in the genome, allowing larger panels, have low level of homoplasy, are easy to standardize with good reproducibility across different laboratories (Zimmerman et al., 2020). SNP genotyping can be performed as panels, also called SNP barcodes, and many platforms have been developed in the last decade for Pv (Baniecki et al., 2015; Kattenberg et al., 2022; Trimarsanto et al., 2022) and Pf (Daniels et al., 2008; Campino et al., 2011; Harrison et al., 2024).

We have developed and validated two amplicon-based next-generation sequencing AmpliSeq assays (Pv AmpliSeq v2 Peru and Pf AmpliSeq v1 Peru assay) for malaria molecular surveillance in Peru (Kattenberg et al., 2023a; Kattenberg et al., 2024). To serve for many uses cases, these assays include molecular resistance and other relevant markers (such as *pfhrp2/3* genes in Pf) and Peru-specific SNP barcodes (composed of 28 SNPs for Pf, and 41 for Pv) for population genetics analysis. These assays enabled to identify predominant Pf lineages with low genetic diversity, double *pfhrp2/3* deletion since 2014 (Kattenberg et al., 2023a), to determine a high heterogeneous Pv transmission with differences according to the settings (Kattenberg et al., 2024), and to study the microepidemiology of an indigenous community with high prevalence and persistence of Pv malaria and a Pf outbreak (Cabrera-Sosa et al., 2024).

As new MMS platforms like the AmpliSeq assays become available, it is relevant to assess whether SNP barcode resolution is comparable to that of the MS panels previously used (Fola et al., 2020; Ghansah et al., 2023). Here, we compared the genetic resolution of the SNP barcodes in the AmpliSeq assays against MS panels to study Pv and Pf populations in the Peruvian Amazon. Specifically, we compared genetic diversity and differentiation metrics and population structure parameters between SNP/AmpliSeq and MS panels. We found that both panels provided comparable results for determining genetic diversity and population structure of Pv and Pf populations in Peru. This information encourages using AmpliSeq assays as an MMS tool to characterize the temporal and spatial dynamics of malaria transmission in Peru. Also important, the comparable results between SNP barcodes and MS allow consistent comparison of population parameters across different studies with either panel. This capability supports their potential implementation into the Peru’s NMEP to enhance monitoring and implement more effective intervention strategies for malaria elimination.

## 2 Materials and Methods

### 2.1 Study samples

Samples used in this study were previously collected from different locations (Supplementary Table S1). First, samples from Nueva Jerusalen (NJ), an indigenous community located northeast of Loreto (region with highest malaria reported cases in Peru), were collected by active case detection (ACD) during November 2019 (Pv, n=48; Pf, n=17) and by passive case detection (PCD) from December 2019 to May 2020 (Pv, n=20; Pf, n=49) (Cabrera-Sosa et al., 2024). In addition, samples from Mazan, a riverine district in Loreto, were collected by two population-based cross-sectional surveys during 2018 (Pv, n=13; Pf, n=10) (Villasis et al., 2021; Rosado et al., 2022). Finally, samples from Santa Emilia (SE), a remote riverine community south of Mazan, were collected by ACD and PCD in 2016 (Pf, n=14). Blood samples (dried filter papers for NJ and SE, or packed red blood cells for Mazan) were collected for molecular diagnosis. Also, thick and thin blood smears were taken for light microscopy diagnosis in the health post of each community. Regardless of the symptoms, treatment was provided for any positive microscopy result, following the national guidelines (Ministerio de Salud, 2015).

### 2.2 Ethics Statement

All sample collections were approved by the Institutional Ethics Committee at Universidad Peruana Cayetano Heredia (UPCH) (SIDISI codes: 64024, 101518 and 102725). Written informed consent was obtained from all participants in Mazan and SE. Samples from NJ were collected as part of the Ministry of Health (MINSA) activities and then transferred to our team for research purposes. This study was also approved by Institutional Ethics Committee at UPCH (SIDISI 207543). All methods were performed following the MINSA guidelines and regulations.

### 2.3 Sample processing

DNA was extracted using EZNA® Blood DNA (Omega Bio-tek, USA), following the manufacturer’s protocol with 50µl as elution volume. Molecular diagnosis was performed by real-time PCR (qPCR). Samples from Mazan were diagnosed using a SYBR Green-based protocol (Mangold et al., 2005). A TaqMan-based assay (Rougemont et al., 2004) was used for samples from NJ and SE.

### 2.4 AmpliSeq assays

Eighty-one Pv and 86 Pf samples with parasitemia greater than 5 parasites/µl were genotyped with the Pv and Pf AmpliSeq Peru assays (Supplementary Fig S1). Library preparation was performed as previously described (Kattenberg et al., 2023a; Kattenberg et al., 2023b; Kattenberg et al., 2024). In brief, 7.5µl of DNA was amplified with two primer pools. Then, digestion, indexes ligation, library amplification and cleaning steps were performed. Libraries were quantified using Qubit High sensitivity DNA kit (Invitrogen), pooled, diluted to 2nM and denatured with NaOH to 7pM. Finally, the denatured library pool was loaded on a MiSeq system (Illumina) for 2x300 paired-end sequencing (Miseq Reagent Kit v3, Illumina) with 5% PhiX spike-in (Illumina).

FASTQ files were processed using an analysis algorithm as previously described (Kattenberg et al., 2023b). After trimming, reads were aligned to the reference genome (PvP01 v46 or Pf3D7 v44 from PlasmoDB, https://plasmodb.org/plasmo/app). Then, variants were called, generating individual gVCF files per each sample, which were combined to call genotypes jointly. After, variants were hard filtered and annotated. Finally, VCFs containing only the Peru SNP barcode in each AmpliSeq assay (28 for Pf, 41 for Pv) were created (Kattenberg et al., 2023a; Kattenberg et al., 2024).

The diploid genotype table (GT) was extracted from the SNP barcode VCF and manually reformatted into a haploid GenAlEx file (Peakall and Smouse, 2012). We constructed multi-locus genotypes (MLG) by combining the alleles across all loci from the barcode and assumed 2 MLG per sample. Then, when a sample had at least one heterozygous position, two MLG were created for GenAlEx file. The reference allele of each heterozygous position was kept in the first “dominant” MLG, and the alternative allele in the other “secondary” MLG, maintaining the homozygous positions in both.

### 2.5 MS Genotyping

In total, 62 Pv and 90 Pf samples were genotyped by using *Plasmodium* species-specific MS panels (Supplementary Fig S1). For Pv, a 16 MS panel (11.162, Ch2.121, 14.297, Ch2.152, Ch14.3021, Ch14.2986, Ch14.3010, Ch2.122, MS9, 13.239, Ch14.2981, MS6, MS4, MS15, 3.502, MS20) previously developed by our team was used (Manrique et al., 2019). Samples were classified according to their parasitemia and followed a different processing before MS genotyping. Samples with >24 molecules/µl (mol/µl) were diluted to 8 - 10 mol/µl for the MS amplification. On the other hand, samples between 4 and 24 mol/µl were amplified by selective whole genome amplification (sWGA) using the enzyme phi29 DNA Polymerase (NEB) before the genotyping. Finally, DNA from samples between 4 and 1.5 mol/µl was reextracted, mixed with the first extraction and used for posterior PCRs (Manrique et al., 2019). Each MS was amplified by conventional PCR. In each reaction, one of the primers was labeled with a fluorophore for later identification. The master mix consisted of 1U of the AccuStart II Taq DNA polymerase enzyme (QuantaBio), 2.5 μL of PCR buffer, 1.5 mM MgCl_2_, 200–400 nM of each primer (depending on the panel) and 2.5 μL of DNA. The amplification protocol consisted of an initial denaturation at 95°C for 5 min, followed by 40 cycles of 30s at 94°C, 30s at 55–67°C (Tm depending on the MS) and 30s at 72°C. The last step was a final extension at 70°C for 15 min (Manrique et al., 2019).

In the case of Pf, a panel of 7 MS (TA1, Poly-α, PFPK2, TA109, 2490, C2M34, C3M69), previously applied to Peruvian samples (Akinyi et al., 2013; Bendezu et al., 2022) was used. Three amplification protocols were used, including two direct PCR (for TA109, Poly-α, PFPK2, C2M34 and C3M69) and one semi-nested PCR (for TA1 and 2490) (Nair et al., 2003; Roper et al., 2003). The master mix consisted of 1X Promega Master-mix (Promega), 400 nM of each primer and 2 µl of DNA or 1 µl of the product of the first PCR. For all cases, extracted DNA was used directly, without dilution or sWGA.

In both Pv and Pf, positive controls and no-template controls were included in the reactions. After MS amplification, the PCR products were diluted and mixed with formamide and the GeneScan 500 LIZ marker (Applied Biosystems). Finally, the mixes were analyzed by capillary electrophoresis on an ABI Prism 3130 (Applied Biosystems). Chromatograms were analyzed with the Microsatellite Analysis software (Applied Biosystems). A quality control of the size marker was performed and the list of peaks per sample was downloaded.

With this data, a semi-automated analysis was performed to select true alleles. First, the peak with the highest relative fluorescence units (RFU) value (height) in the no-template controls was used as a threshold and all peaks with lower height in samples were discarded. After, the peak with the highest height was considered the main allele for each MS and sample. Secondary alleles were considered if their peak height was at least 1/3 of the main allele, and their size was not within ± 2bp of the main allele. As a final validation step, we manually compared the selected alleles in each sample to the positive control, discarding any allele when a discrepancy in the shape or pattern of peaks was detected. If a previously selected main allele was discarded in this process, the remained allele with the highest height was reclassified as the new main allele.

The final allele table was reformatted manually into a haploid GenAlEx file (Peakall and Smouse, 2012). For subset analysis, we separated the data of samples with secondary alleles into 2 MLG in the GenAlEx formatted file, with the main alleles kept in the first “dominant” MLG, and the secondary alleles in the other MLG, maintaining the rest of MS alleles in both.

### 2.6 Sample and makers quality control

We applied some inclusion criteria for the population genetics analyses (Supplementary Fig S1). First, to ensure comparability, we selected samples with genotyping data from both AmpliSeq and MS (Pv, n = 62; Pf, n = 86). Then, samples with missing data in more than 25% of panels (SNP or MS) were excluded. 80/86 Pf and 51/62 Pv samples passed this filter. Next, we kept markers with less than 25% missing data. Using this criteria, 2/28 positions from the Pf SNP barcorde and 1/16 MS from the Pv panel were excluded. At the end, the final genotyped data included 80 Pf samples with 7 MS (105 MLG) and 26 SNP (117 MLG), and 51 Pv samples with 15 MS (86 MLG) and 40 SNP (68 MLG) (Supplementary File 1).

### 2.7 Data and population genetic analysis

We assessed the inclusion of the secondary MLG in the analysis by comparing the genetic differentiation of MLG sets containing: Set 1) samples with only one MLG, Set 2) dominant MLG of all samples, Set 3) dominant MLG of all samples and secondary MLG of samples with in only one secondary allele and Set 4) all dominant and secondary MLG of all samples. Fixation index (Fst) was calculated using hierfstat package in R (Goudet et al., 2022). Genetic differentiation (Fst) between all these sets were close to 0 (Supplementary Fig. S2), indicating no genetic differences between them. Therefore, all MLG for all samples were included.

We calculated population genetic parameters using each SNP and MS panel for Pv and Pf samples. Genetic diversity was measured as expected heterozygosity (He) and calculated using the formula: He = [n/(n − 1)][1 − Σp2], where n is the number of genotyped samples and p is the frequency of each allele at a given locus (Nei and Roychoudhury, 1974; Serrote et al., 2020). Genetic differentiation was expressed as pairwise Fst values and 95% confidence intervals (95% CI), calculated based on Weir and Cockerham’s Method using the diveRsity package in R (Weir and Cockerham, 1984; Keenan et al., 2013). Population structure was explored using principal component analysis (PCA) (Jolliffe and Cadima, 2016) and the software STRUCTURE v2.3 (Pritchard et al., 2000). PCA was performed using the “prcomp” function in stats R-package. STRUCTURE determined the most likely number of clusters (K). Runs were performed exploring K from 1 to 10 (20 iterations each), assuming a mixture model, correlated allele frequencies, using sampling location as priors (LOCPRIOR option). They had a burn-in period of 50,000 iterations followed by 150,000 Markov Chain Monte Carlo (MCMC) iterations. The most likely K was defined by calculating the rate of change of K (ΔK) (Evanno et al., 2005), using the R package pophelper (Francis, 2017). The complexity of infection (COI) was used to assess the proportion of polyclonal infections (multiple clones in one sample). Samples with 2 MLG (i.e., at least one heterozygous SNP genotype for the AmpliSeq data or at least one secondary allele in any MS) were considered polyclonal. For comparison, we also determined COI with the within-sample F (Fws) statistic using all biallelic SNPs detected by the AmpliSeq assays (Pv: 2923 SNPs, Pf: 963 SNPs) using the moimix package in R. Samples were considered a monoclonal infection when the Fws was ≥ 0.95 (Auburn et al., 2012).

### 2.8 Statistical analysis

He values across different locations were compared performing the Kruskal-Wallis rank sum test. Dunn’s post hoc tests were used to perform pairwise comparisons between all possible pairs of groups to determine which specific groups are significantly different from each other. Comparison of He results obtained with MS and SNP were calculated using the correlation coefficient using Spearman’s rank correlation. Pairwise Fst from SNP and MS data were compared using Wilcoxon Signed-Rank Test. Proportions of mono/polyclonal infections according to SNP barcode, MS and Fws were compared for each species (Pf and Pv) using the chi-squared test. Cohen’s Kappa coefficient was used to assess the agreement between SNP and MS in determining the complexity of infection. All statistical analyses were performed in R Studio (version 2022.12.0). P-values < 0.05 were considered significant. The Bonferroni correction was applied to adjust the p-values when performing multiple pairwise comparisons

### 2.9 Cost estimates

Costs associated with AmpliSeq and MS genotyping for 96 samples (and controls) were estimated, including materials (reagents and kits) and time-personnel costs. All material prices were calculated based on quotations and invoices from Lima, Peru, from 2022. The cost does not consider DNA extraction, molecular diagnosis and plastic labware because those methods are similar for both procedures. For Pv MS genotyping, we assumed that 60% of samples required sWGA (Manrique et al., 2019). We did not add the cost of sequencing and capillary electrophoresis as we assumed the necessary equipment, such as the MiSeq and ABI Prism 3130, was already available and maintained. Total costs included portion of materials used exactly for 96 samples.

Personnel cost per day (60 USD/working day) was calculated considered monthly salary of 1200 USD and 20 working days in 1 month. Number of working days in each activity was based on practical experience. Costs in Peruvian Nuevos Soles (PEN) were converted into United States Dollars (USD) using the exchange rate of 1 PEN = 0.26 USD.

## 3 Results

### 3.1 AmpliSeq and MS genotyping performance

SNP barcode performances in the Pv and Pf AmpliSeq assays in these samples were previously described (Cabrera-Sosa et al., 2024; Kattenberg et al., 2024). For Pv, only the reference allele was detected for the SNP PvP01_13_v1_32509. The median minor allele frequency (MAF) for the rest of 40 SNPs was 0.27 (range: 0.02-0.46, mean ± standard deviation (sd): 0.25 ± 0.14). All Pv samples had ≤25% missing genotypes in the barcode. In the case of Pf, all alternate alleles at 28 SNPs were genotyped successfully. The median MAF was 0.19 (range: 0-0.48, mean ± sd: 0.20 ± 0.15). Two SNPs (Pf3D7_02_v3_694307 and Pf3D7_09_v3_231065) had >25% of missing data (57 and 66%, respectively) and were excluded for posterior analyses. A low proportion of samples (4.65%, 4/86) had >25% of missingness in the SNP barcode and were subsequently excluded.

The amplification efficiency of both MS panels is shown in Supplementary Figure S3. For Pv, MS had low proportion of missing data (3.2 – 22.6%), except for Ch14.3010 (43.6%), which was excluded for later analyses. 52 (83.4%) samples had ≤ 25% missing genotypes. On the other hand, amplification efficiency in the Pf MS panel was ≥90% (% missing data: 0 – 10%). Only 5 samples (5.6%) with >25% of missingness were excluded.

### 3.2 Genetic diversity

We found similar trends of population diversity (expressed as He) with both panels (MS or SNP) for both species (Fig. 1, Supplementary Figure S4). Pv populations in Mazan and NJ had similar genetic diversity when using MS (Fig. 1A, median He: 0.68 – 0.73, p = 0.23) or SNP (Fig. 1B, median He: 0.37 – 0.38, p = 0.80). We also noted that 2/40 (5%) of the SNP positions were fixed in all population (Supplementary Table S2).

**Figure 1.**
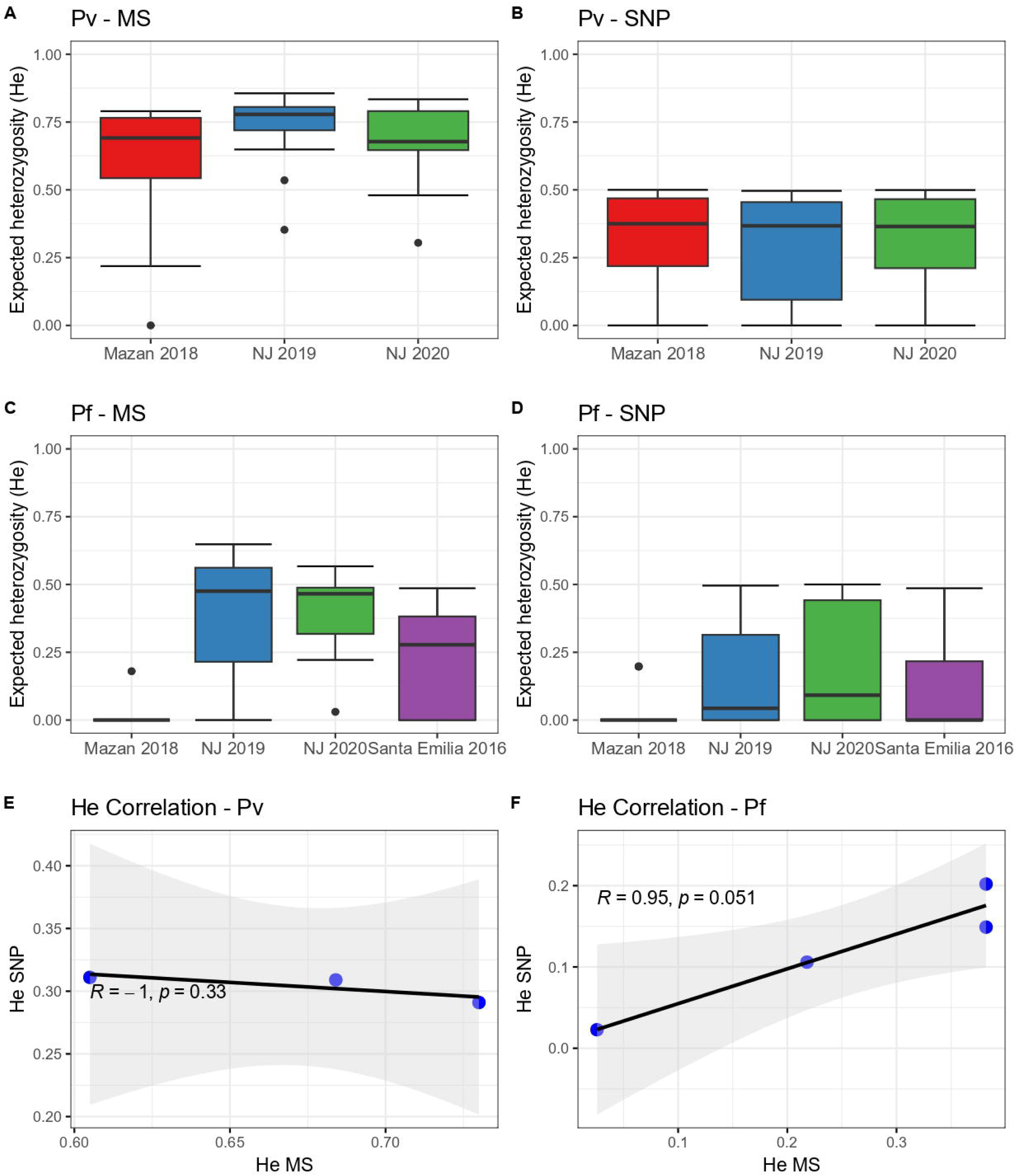
Expected heterozygosity (He) in Pv and Pf populations using SNP and MS. He was calculated for Pv (A and B) and Pf (C and D) using MS (A and C) or SNP (B and D). The horizontal line within the boxes represents the median He from each panel (MS) or position (SNP) for all samples in each group. Dots outside the boxes represent the outlier data. In addition, Spearman’s correlations of He values by MS and the SNP barcode for (E) Pv and (F) Pf populations were determined.

On the other hand, Pf isolates from Mazan (median He: 0 with both markers) had lower diversity than parasites from NJ using MS (Fig. 1C, median He: 0.36 – 0.37, p = 0.03 – 0.04) or SNP (Fig. 1D, median He: 0.04 – 0.09, p = 0 – 0.03). Also, He values in Pf samples from Mazan and NJ were similar to Santa Emilia isolates, determined by MS (median He: 0.28, p = 0.78 – 1) and by SNP (median He: 0, p = 0.36 – 1). Finally, around 40% of SNP positions (10/26) were fixed in these populations (Supplementary Table S2).

Despite the similar trends in the genetic diversity, we found no significant correlation between He values obtained by MS and SNP in the Pv (Fig. 1E, ρ = -1.00, p = 0.33) and Pf (Fig. 1F, ρ = 0.95, p = 0.051) populations, although the correlation lacked statistical power as the degrees of freedom were low (1 and 2, respectively).

### 3.3 Genetic differentiation

We observed no significant differences between the pairwise Fst values obtained using MS and SNP across the Pv (p = 0.34 – 0.85) and Pf (p = 0.24 – 0.83) populations (Fig. 2). In case of Pv, low differentiation of parasites within NJ was noted when using MS (Fst = 0.04, 95% CI: 0.01 – 0.07) or SNP (Fst = 0.03, 95% CI: 0 – 0.09) (Fig. 2A). Moderate levels of differentiation were observed between Mazan and NJ with MS (Fst = 0.13 – 0.14, 95% CI: 0.08 – 0.22) and SNP (Fst = 0.09 – 0.12, 95% CI: 0.02 – 0.22) (Fig. 2A). In the Pf population, the lowest Fst values were found between the two groups in NJ when using MS (Fst = 0.14, 95% CI: 0.07 – 0.24) and SNP (Fst = 0.19, 95% CI: 0.09 – 0.32). On the other hand, the highest Fst values was observed between Mazan and SE isolates (Fst = 0.65, 95% CI: 0.41 – 0.84 with MS, Fst = 0.61, 95% CI: 0.31 – 0.83 with SNP) (Fig. 2B). The consistency between the two markers suggest that both MS and SNP markers are capable of reliably capturing genetic differentiation in the Peruvian setting.

**Figure 2.**
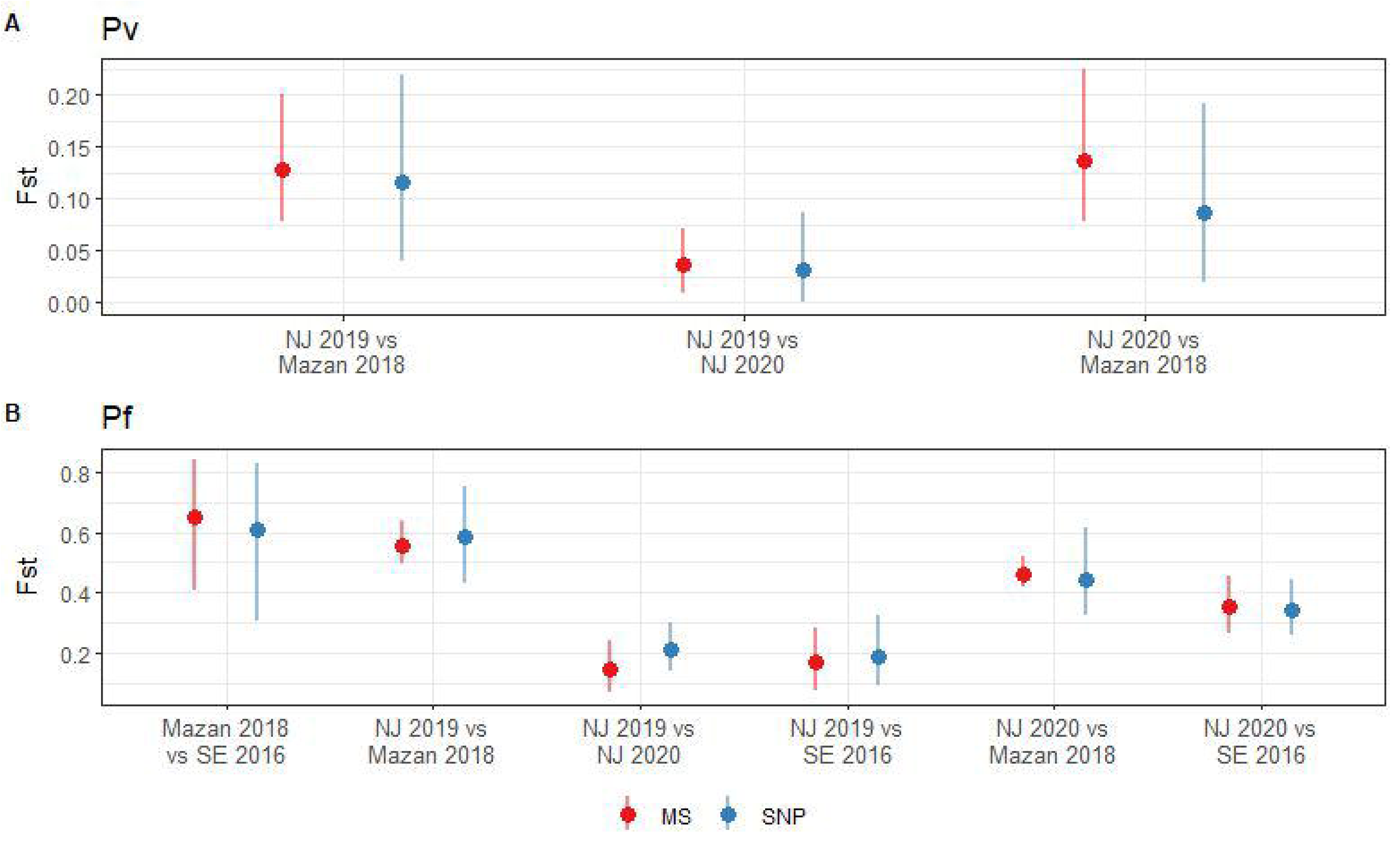
Pairwise fixation index (Fst) in Pv and Pf populations using SNP and MS. Fst was calculated for Pv (A) and Pf (B) using MS (red) or SNP barcode (blue). Pairwise Fst values (points) and 95% confidence intervals (lines) were estimated with 1000 bootstraps using the diveRsity package in R.

### 3.4 Population structure

Clustering patterns obtained from STRUCTURE were strikingly similar between MS and SNP in both species (Fig. 3), although the most likely number of clusters (K) differed between panels. For Pv, the predicted best K were 2, 4 and 6 according to the SNP barcode, whereas the best K were 3 and 6 using MS (Supplementary Fig. S5). For Pf, the predicted best K using SNP were 3, 5 and 7, meanwhile the most likely K with MS were 2, 4 and 7 (Supplementary Fig. S6). For comparison between MS and SNP, we plotted the assignation to 3, 4 and 6 clusters for Pv isolates (Fig. 3A and Supplementary Fig S5) and to 4, 5 and 7 clusters for Pf parasites (Fig. 3B and Supplementary Fig. S6).

**Figure 3.**
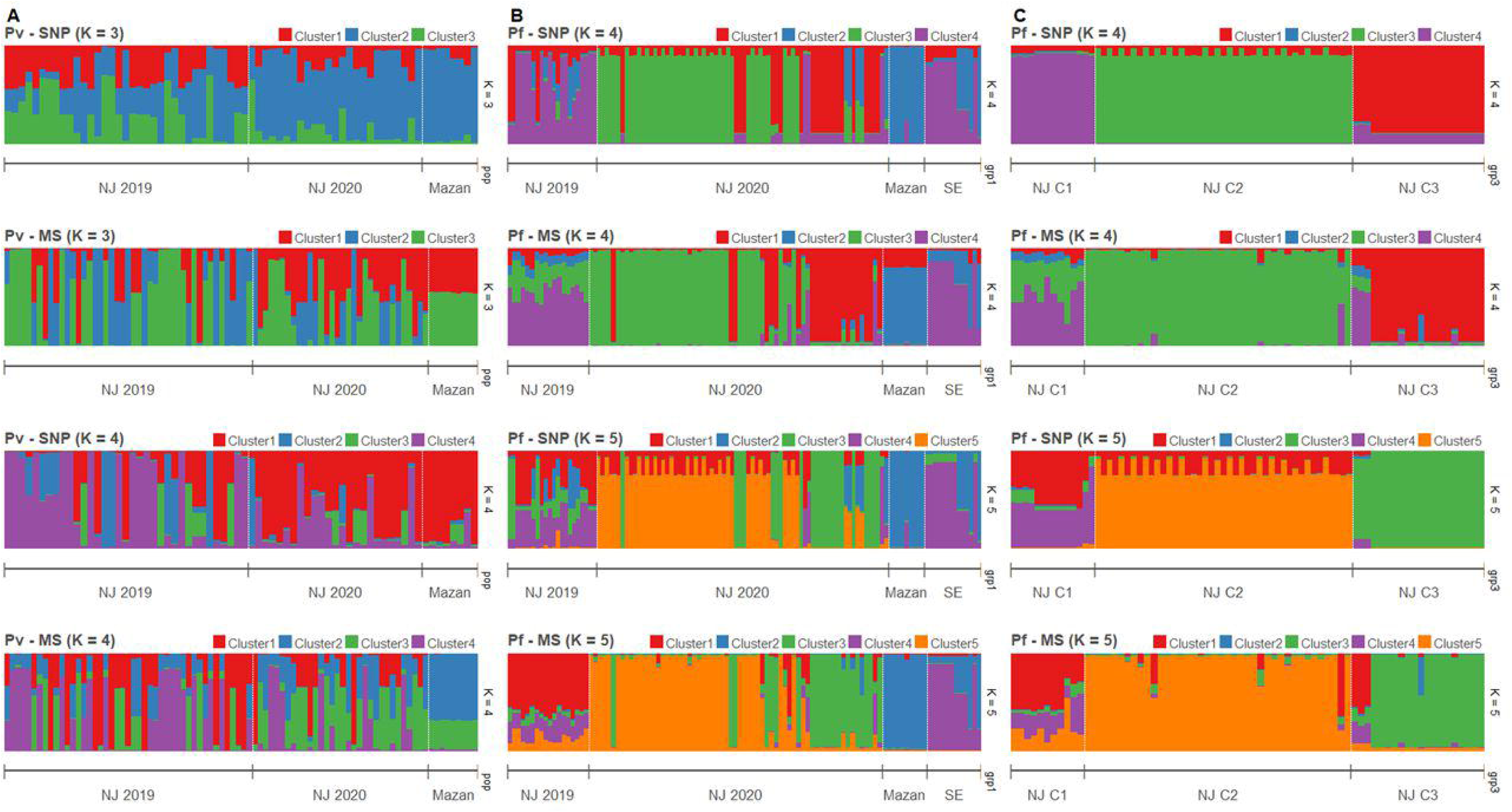
Clustering analysis by STRUCTURE in Pv and Pf populations using SNP and MS. The graph depicts the clustering models when the isolates were assigned into different number of clusters (K). Each isolate is represented by a single vertical line broken into K coloured segments, with lengths proportional to each of the K inferred clusters. **(A)** Clustering of Pv populations sorted by location at K = 3 and 4 when using SNP and MS. **(B)** Clustering of Pf populations sorted by location at K = 4 and 5 when using SNP and MS. **(C)** Clustering of Pf populations in NJ by previous classification according to (Cabrera-Sosa et al., 2024) at K = 4 and 5 when using SNP and MS.

For Pv populations, high admixture was observed in NJ, with all clusters present in both years. In contrast, all samples from Mazan belongs to predominantly one cluster according to SNPs, while MS did not attribute to any single cluster but showed the same admixture pattern. (Fig. 3A, Supplementary Fig. S5).

In the case of Pf, strong clustering observed in all population, except in NJ 2019, which presented high admixture. Predominant clusters in NJ 2020 and Mazan were different, and SE shared some clusters with NJ 2019 and Mazan. SNPs also detected some admixed samples in NJ 2020, which are predicted to belong to a single cluster by MS (Fig. 3B and Supplementary Fig. S5).

Previously, we identified three genetic clusters (C1 to C3) in the Pf population in NJ using all biallelic SNPs detected in the AmpliSeq assay based on results from PCA and supported by other analyzes such discriminant analysis of principal components (DAPC), identity by descent (IBD) networkand phylogenetics (Cabrera-Sosa et al., 2024). Using STRUCTURE analysis, all those genetic cluster had distinct patterns with only the 28-SNP barcode or MS (Fig. 3C and Supplementary Fig. S6). In C1, similar admixture clustering was frequent, except when K = 4 with SNPs were plotted. Different clusters were predominant in each C2 and C3. (Fig. 3C and Supplementary Fig. S6).

PCA analysis confirmed patterns observed in the STRUCTURE analysis, and overall similar (lack of) clustering by both MS and SNP (Fig. 4). For Pv, no clear geographical or temporal clustering was observed in neither panel (Fig. 4A and 4B). On the other hand, a stronger clustering of Pf isolates of NJ 2020 was observed in SNP than with MS (Fig. 4C and 4D). The variance explained by the first two principal components were similar for Pv (20% and 25% for MS and SNP, respectively) and Pf (63% for MS, 62% for SNP).

**Figure 4.**
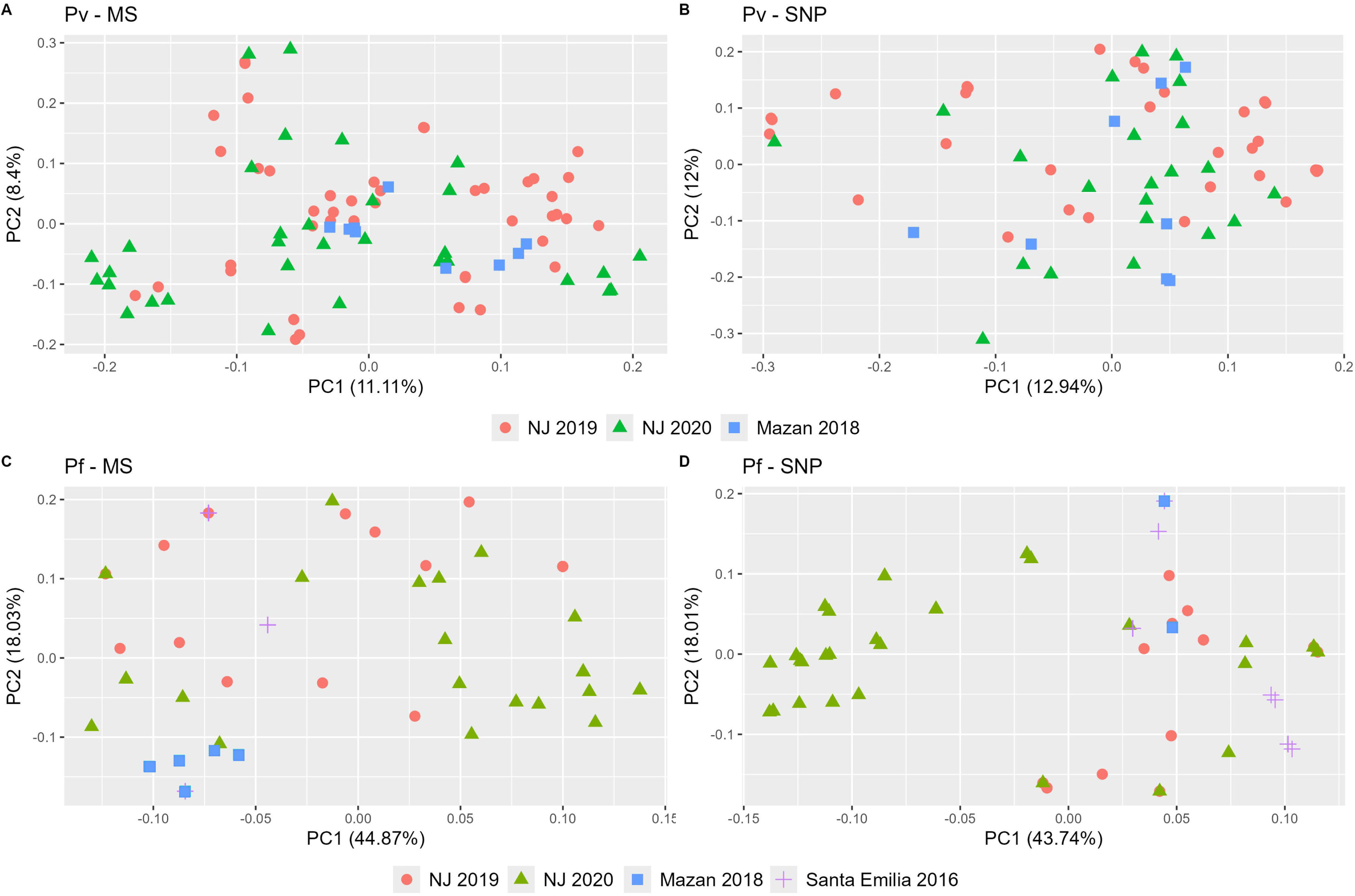
Principal component analysis (PCA) of Pv and Pf populations using SNP and MS. PCA was performed for Pv (A and B) and Pf (C and D) using MS (A and C) or SNP (B and D). Each community is represented by a unique shape and color. Consistent shapes are used to identify the same community across both Pv and Pf populations.

### 3.5 Complexity of Infection (COI)

We found differences in the proportion of polyclonal infections determined by the SNP barcode, MS and the Fws statistics (Table 1). For Pv, this proportion was higher with MS (69%) than with SNP and Fws (33% and 28%, respectively; p = 3.3x10^-5^). Five MS (MS9, MS15, Ch2.121, Ch2.122, Ch2.152) accounted for high proportion of samples with secondary alleles (26 – 43%). For Pf, the SNP barcode detected similar polyclonal infections as MS (46.2 and 31.2%, respectively, p = 0.21), but higher than Fws (29%, p = 0.043). Agreement between SNP barcode and MS was higher in PF (κ = 0.54, 95% CI: 0.36 – 0.72) than in Pv (κ = 0.23, 95% CI: 0.04 – 0.43).

**Table 1.** Proportion of polyclonal infections by SNP barcode, MS and Fws in Pv and Pf samples. Samples were considered polyclonal with at least one heterozygous position (SNP barcode), secondary allele (MS) or Fws < 0.95.

| Population | Pv (n = 51) |  |  |  | Pf (n = 80) |  |  |  |
| --- | --- | --- | --- | --- | --- | --- | --- | --- |
|  | SNP<br>barcode -<br>AmpliSeq | MS | Fws | p-value | SNP<br>barcode -<br>AmpliSeq | MS | Fws | p-value |
| NJ 2019 | 8 (29.6%) | 18<br>(66.7%) | 6<br>(22.2%) | **$3.3 \times 10^{-5}$** | 8 (57.1%) | 4<br>(26.8%) | 3<br>(20.1%) | **0.01** |
| NJ 2020 | 7 (38.9%) | 14<br>(77.8%) | 6<br>(33.3%) |  | 27 (60%) | 20<br>(44.4%) | 19<br>(42.2%) |  |
| Mazan | 2 (22.2%) | 3<br>(33.3%) | 2<br>(22.2%) |  | 0 (0%) | 1<br>(11.1%) | 0 (0%) |  |
| SE |  |  |  |  | 2 (16.7%) | 0 (0%) | 1<br>(8.4%) |  |
| **Total** | **17**<br>**(33.3%)** | **35**<br>**(68.6%)** | **14**<br>**(27.5%)** |  | **37**<br>**(46.2%)** | **25**<br>**(31.2%)** | **23**<br>**(28.8%)** |  |

### 3.6 Cost estimations

We estimated higher costs associated with AmpliSeq compared to MS genotyping (Table 2). Th estimated cost of the AmpliSeq was 183 USD/per sample, whereas using the MS panels was 49 USD/sample for Pv (16 MS) and 27 USD/sample for Pf (7 MS). Notably, a large difference was observed for the personnel costs, with the AmpliSeq been 2 to 4 times cheaper than MS genotyping (5 vs 11 – 19 USD/sample), as it was much faster (AmpliSeq: 6 working days, MS: 18 to 30 working days) (Supplementary File 2).

**Table 2.** Estimated costs of AmpliSeq and MS genotyping for 96 samples.

|  | Pv or Pf AmpliSeq |  |  | Pv MS (16 MS) |  |  | Pf MS (7 MS) |  |  |
| --- | --- | --- | --- | --- | --- | --- | --- | --- | --- |
|  | Total<br>cost<br>(PEN) | Total<br>Cost<br>(USD) | Cost/<br>sample<br>(USD) | Total<br>cost<br>(PEN) | Total<br>Cost<br>(USD) | Cost/<br>sample<br>(USD) | Total<br>cost<br>(PEN) | Total<br>Cost<br>(USD) | Cost/<br>sample<br>(USD) |
| Materials | 65695 | 17081 | 118 | 11253 | 2926 | 30 | 5800 | 1508 | 16 |
| Personnel | 1846 | 480 | 5 | 6923 | 1800 | 19 | 4154 | 1080 | 11 |
| **Total** | **67541** | **17561** | **183** | **18176** | **4726** | **49** | **9954** | **2588** | **27** |

## 4 Discussion

The use of genomic tools for malaria parasite genomic surveillance has increased over the last decade. In the past, genetic panels such as MS were mainly used to characterize genetic population of malaria parasites, particularly using panels composed of relatively few MS, typically <20 (Sutton, 2013; Manrique et al., 2019; Gwarinda et al., 2021). More recently, the technological advancements in DNA sequencing enable researchers to analyze many loci at once. Due to their higher prevalence in the genome, potential to target functional regions, and the fact that MS regions often perform poorly in targeted sequencing (Rovira-Vallbona et al., 2023b), SNPs have been replacing MS in population genetic studies (Zimmerman et al., 2020; Neafsey et al., 2021). As new technologies involving SNP barcodes for surveillance appear, a comparison with MS genotyping could offer valuable insights into the similarity of output results and the limitations and advantages of each method and their applications under different scenarios. In this study, we demonstrated good agreement between SNP barcodes from our novel AmpliSeq assays (Kattenberg et al., 2023a; Kattenberg et al., 2024) and MS panels for population genetic analyses in the same group of Pv and Pf samples from the Peruvian Amazon, showing that similar conclusions on the transmission dynamics can be concluded with both panels.

In the case of the diversity metrics, both panels yielded relatively consistent trends. For Pv, He values were consistent across the different study settings in MS and SNP results. For Pf, both panels agreed on the diversity differences between Mazan and NJ. This observation shows that the SNP barcode and MS could capture a high portion of the genome-wide genetic similarly, even comparable to larger panels such > 2000 biallelic SNPs (Kattenberg et al., 2024). This is also consistent to previous studies about comparison of SNP and MS diversity estimates in other species (Zimmerman et al., 2020; Perez-Gonzalez et al., 2023). As expected, He values derived from MS were higher across all populations for both Pv and Pf populations compared to SNP, explained by the higher theoretical maximum heterozygosity in each panel (He = 1 for MS vs He = 0.5 for SNP) (Zimmerman et al., 2020). In both panels, Pv population was more diverse than Pf in the Peruvian Amazon, and association of genetic diversity with transmission intensity (NJ > Mazan) were observed in Pf, but not in Pv, all consistent with reports in other locations (Barry et al., 2015; Auburn et al., 2021). In that sense, our results are crucial because genetic diversity can be reliably estimated with SNP or MS and the similar trends allow to compare data from different studies that have used those panels.

Generally speaking, genetic differentiation estimates (Fst) from MS panels and the SNP barcode were similar in the Pv and Pf populations. Except for the Pv isolates within NJ, most of the pairwise comparisons showed moderate or great differentiation with both MS and SNP. This consistency between the both panels for the genetic differentiation in *Plasmodium* was observed in other species (Zimmerman et al., 2020; Perez-Gonzalez et al., 2023), showing that the level of differentiation can be reliable determined using either MS panel or the SNP barcode. This measure provides valuable insights into the population structure, helps estimate gene flow among parasite populations, and can identify potential parasite migration route between locations, which are correlated to human mobility in most of cases (Escalante et al., 2015; Escalante and Pacheco, 2019; Tessema et al., 2019), although other factors such as migration rates and population sizes may also contribute to the genetic differentiation (Meirmans and Hedrick, 2011; Kitada et al., 2021)

The population structure analyses demonstrated that although given similar clustering patterns, the SNP barcode provided greater precision, resulting in more distinct grouped individuals than the more loosely clustered individuals observed with MS panels, particularly for Pf. In that sense, we found different optimal number of clusters when using MS and SNP in Pv and Pf, which is in concordance to other reports in mammals, reptiles and amphibians (Camacho-Sanchez et al., 2020; Biello et al., 2022; Perez-Gonzalez et al., 2023). This variation can be explained by number of samples per population and number of loci, which strongly affect the K determination and population assignment. Also, as SNPs have fewer possible variations, a change in one of their alleles might have a bigger impact on the results, making it more efficient to detect clusters than with a change of an alleles in the larger MS pool (Puckett and Eggert, 2016; Camacho-Sanchez et al., 2020). Regardless, both MS and the SNP barcode could detect and differentiate the reported genetic clusters in the Pf population in NJ (Cabrera-Sosa et al., 2024), making both panels suitable for identifying outbreaks. Once again, these findings suggest that both panels can be effectively used to investigate population structure.

The contrasting results between SNP barcode, MS and Fws show the difficulties involving COI determination (Zhong et al., 2018). For Pv, the proportion of polyclonal infections was higher with the MS panel than the SNP barcode, while for Pf, the proportions were similar between both panels, but SNP barcode results were higher than calculations with Fws. Previous reports have shown that SNP barcodes may underestimate COI compared to MS in Pf populations in Vietnam and Peru (Kattenberg et al., 2023a; Rovira-Vallbona et al., 2023a), in contrast to a correlation between MS and Fws outputs for COI in a Pf population from Guinea (Murray et al., 2016). MS could detect additional genotypes than SNP as their higher mutation rate leads to more alleles and more information content per locus (Sutton, 2013; Puckett and Eggert, 2016). Although higher with MS, proportion of polyclonality determined by SNP barcode in the Pv population was similar to previous reports in the Peruvian Amazon (Delgado-Ratto et al., 2016; Manrique et al., 2019; Kattenberg et al., 2024). However, factors such as the time and location should be considered as could showed differences in the transmission intensity. In that regard, several studies have demonstrated the correlation between COI and prevalence or transmission level in Pv and Pf populations around the world (Escalante et al., 2015; Fola et al., 2017; Kattenberg et al., 2020; Lopez and Koepfli, 2021), with some exceptions mainly observed in Pv (Escalante and Pacheco, 2019; Auburn et al., 2021). In our study, the Pv transmission levels in NJ (high) and Mazan (low) correlated with their proportion of polyclonal infections. This association was not observed with the genetic diversity, showing that COI may be used as a more effective proxy for transmission intensity. As the SNP barcodes were not specifically designed for measuring COI, MS panel’s allelic richness, particularly of MS9, MS15, Ch2.121, Ch2.122 and Ch2.152, make them more suitable for COI determination’s studies in Pv populations (Sutton, 2013). Finally, other markers (such as *ama1*, *csp* and *cpmp* genes) can be used to determine COI in *Plasmodium* populations (Zhong et al., 2018; Wamae et al., 2022). As the *ama1* gene is targeted in our AmpliSeq assays, this marker may complement the SNP barcode to assess COI (Kattenberg et al., 2023a; Kattenberg et al., 2024).

Previous studies have compared both panels (SNP and MS) in *Plasmodium* populations, reporting mixed results. SNPs have been reported to have better resolution than MS in Pv isolates from Papua New Guinea in 2012–14 (Fola et al., 2020). On the other hand, SNPs and MS were able to correctly cluster Pf from Africa, but only SNPs correctly classified strains from South and Central America (Kanai et al., 2022). In contrast, SNPs had lower genetic resolution to identify changes after an intervention in a Pf population in Ghana (Ghansah et al., 2023). Finally, similar genetic differentiation and population structured were observed with both SNP and MS in Pf populations from Western Keyna (Lo et al., 2018). In our study, comparable results were obtained with SNPs and MS in Pf and Pv populations in Peru, with slightly higher resolution using the first in some analysis, which allow to compare evolution of population genetics parameters through space and time. The differences in the geographic scale may be one explanation of the differences to the previous reports. Here, we worked at a small geographic scale, which is required when malaria elimination scenario is close (Auburn et al., 2021). Another explanation is the design of the SNP barcodes, which was based on whole genomic available data (Kattenberg et al., 2023a; Kattenberg et al., 2024). As also previously reported, we showed some fixed SNP positions, particularly in the Pf population, that may affect how much diversity the barcode can capture. Now, with higher available up to date genomes from the region, the resolution of the SNP barcode could be increased. Other factors can include number of panels and sample size, different mutation rates, diverse transmission level, and changes in the informativeness over time and place, making necessary to validate the panels in new scenarios (Delgado-Ratto et al., 2016; Manrique et al., 2019; Taylor et al., 2019).

Our findings highlighted that comparing the efficiency of SNP barcodes in the AmpliSeq assays and MS panels for malaria parasite population genetics is complex and varies depending on the type of analysis and the specific parasite species. Even if SNP and MS panels remain comparable in different scenarios, each panel has challenges for their implementation into NMCP/NMEPs that should be considered (Ruybal-Pesantez et al., 2024). MS are highly polymorphic and have low cost. However, their genotyping is difficult to reproduce and standardize between different laboratories, and involves long times for laboratory and analysis work and may be not suitable for larger sample sets and timely results required for MMS (Puckett and Eggert, 2016; Zimmerman et al., 2020). Also, results may be affected by subjectivity during allele determination, requiring automated methods to avoid these issues (Manrique et al., 2015; Barbian et al., 2018; Salado et al., 2021). On the other hand, key benefits of the AmpliSeq assays are their suitability for high-throughput and automated analysis, facilitating the rapid generation of big amount of genetic information. While notably more expensive than MS, these assays also include more than 10 drug resistance associated genes panels and *pfhrp2/3* genes (Kattenberg et al., 2023a; Cabrera-Sosa et al., 2024; Kattenberg et al., 2024). Also, they become cheaper with more processed samples (Supplementary File 2). Currently, we are testing other reagents with the same panel to reduce the library preparation costs. However, they required specialized technology and bioinformatic expertise skills, not always particularly accessible in remote areas of low and middle-income countries. This requires a better distribution of sequencing infrastructure and priority efforts from the health authorities, as seen with other pathogens in Peru (Padilla-Rojas et al., 2021; Puyen et al., 2024).

In conclusion, cost-effective and scalable platforms should be incorporated into NMCP/NMEPs for molecular surveillance. Here, we showed that the SNP barcode in the Pv and Pf AmpliSeq assays provide similar results to classic MS genotyping in population genetics analyses such as patterns of transmission intensity and population structure. As both types of panels are comparable, the selection of the best option should be based on the objectives and practical limitations, including cost, available infrastructure, feasibility, and the specific research questions (Kanai et al., 2022). In any case, while tackling the associated challenges, the implementation of novel NGS tools, such the AmpliSeq assays, into the surveillance system in Peru may be encouraged.

## 7 Author Contributions

Conceptualization: LCS, CDR. Data curation: LCS, MS. Formal analysis: LCS, MS. Funding acquisition: CDR, DG, ARU, JV, LCS. Investigation: LCS, MS, RR. Resources: JV, ARU, DG. Supervision: CDR. Validation: LCS, JHK, ARU, CDR. Visualization: LCS, MS. Writing – original draft: LCS, MS, CDR. Writing – review & editing: LCS, MS, CDR, ARU, JHK. All authors approved the final version of the manuscript

## 8 Funding

This work was funded by VLIR-UOS (PE2018TEA470A102), the Research Foundation-Flanders (FWO, G.0A42.22N), Belgium Development Cooperation (DGD) under the Framework Agreement Program FA4 Peru (2017–2021) and FA5 Peru (2022–2026) and a doctoral scholarship from the National Council for Science, Technology, and Technological Innovation (CONCYTEC) Peru, through its executing unit National Fund for the Development of Science, Technology, and Technological Innovation (FONDECYT) Perú contract N° 165-2020-FONDECYT to LCS. Sample collection in Mazan and SE was supported by NIH (5U19AI089681-15). The funders had no role in study design, data collection, analysis, publication decision, or manuscript preparation.

### 9 Acknowledgments

First, we want to thank all the participants and workers of the projects included in this study. We acknowledge Prof. Jean-Pierre Van geertruyden (Global Health Institute), Dr. Hugo Valdivia (Naval Medical Research Unit SOUTH) and Prof. Dr. Alejandro Llanos-Cuentas (Instituto de Medicina Tropical "Alexander von Humboldt") for their contribution on NJ and SE data acquisition. Finally, we want to thank Mr. Jhon Zumaeta for his help in performing the STRUCTURE runs.

## 10 Data Availability Statement

Pv and Pf sample metadata and allele tables (in GenAlEx format) are available in Supplementary File 1. Raw data (FASTQ files) from the AmpliSeq assays are available at the SRA under BioProject accession numbers PRJNA1055117 (Pv) and PRJNA1074830 (Pf). Scripts are available upon request. All other data are included in the manuscript and supporting information.

## References

Akinyi, S., Hayden, T., Gamboa, D., Torres, K., Bendezu, J., Abdallah, J.F., et al. (2013). Multiple genetic origins of histidine-rich protein 2 gene deletion in Plasmodium falciparum parasites from Peru. Sci Rep 3, 2797. doi: 10.1038/srep02797.

Anderson, T.J., Haubold, B., Williams, J.T., Estrada-Franco, J.G., Richardson, L., Mollinedo, R., et al. (2000). Microsatellite markers reveal a spectrum of population structures in the malaria parasite Plasmodium falciparum. Mol Biol Evol 17(10), 1467–1482. doi: 10.1093/oxfordjournals.molbev.a026247.

Auburn, S., Campino, S., Miotto, O., Djimde, A.A., Zongo, I., Manske, M., et al. (2012). Characterization of within-host Plasmodium falciparum diversity using next-generation sequence data. PLoS One 7(2), e32891. doi: 10.1371/journal.pone.0032891.

Auburn, S., Cheng, Q., Marfurt, J., and Price, R.N. (2021). The changing epidemiology of Plasmodium vivax: Insights from conventional and novel surveillance tools. PLoS Med 18(4), e1003560. doi: 10.1371/journal.pmed.1003560.

Baniecki, M.L., Faust, A.L., Schaffner, S.F., Park, D.J., Galinsky, K., Daniels, R.F., et al. (2015). Development of a single nucleotide polymorphism barcode to genotype Plasmodium vivax infections. PLoS Negl Trop Dis 9(3), e0003539. doi: 10.1371/journal.pntd.0003539.

Barbian, H.J., Connell, A.J., Avitto, A.N., Russell, R.M., Smith, A.G., Gundlapally, M.S., et al. (2018). CHIIMP: An automated high-throughput microsatellite genotyping platform reveals greater allelic diversity in wild chimpanzees. Ecol Evol 8(16), 7946–7963. doi: 10.1002/ece3.4302.

Barry, A.E., Waltmann, A., Koepfli, C., Barnadas, C., and Mueller, I. (2015). Uncovering the transmission dynamics of Plasmodium vivax using population genetics. Pathog Glob Health 109(3), 142–152. doi: 10.1179/2047773215Y.0000000012.

Bendezu, J., Torres, K., Villasis, E., Incardona, S., Bell, D., Vinetz, J., et al. (2022). Geographical distribution and genetic characterization of pfhrp2 negative Plasmodium falciparum parasites in the Peruvian Amazon. PLoS One 17(11), e0273872. doi: 10.1371/journal.pone.0273872.

Biello, R., Zampiglia, M., Fuselli, S., Fabbri, G., Bisconti, R., Chiocchio, A., et al. (2022). From STRs to SNPs via ddRAD-seq: Geographic assignment of confiscated tortoises at reduced costs. Evol Appl 15(9), 1344–1359. doi: 10.1111/eva.13431.

Brashear, A.M., and Cui, L. (2022). Population genomics in neglected malaria parasites. Front Microbiol 13, 984394. doi: 10.3389/fmicb.2022.984394.

Brown, T.S., Arogbokun, O., Buckee, C.O., and Chang, H.H. (2021). Distinguishing gene flow between malaria parasite populations. PLoS Genet 17(12), e1009335. doi: 10.1371/journal.pgen.1009335.

Cabrera-Sosa, L., Nolasco, O., Kattenberg, J.H., Fernandez-Minope, C., Valdivia, H.O., Barazorda, K., et al. (2024). Genomic surveillance of malaria parasites in an indigenous community in the Peruvian Amazon. Sci Rep 14(1), 16291. doi: 10.1038/s41598-024-66925-x.

Camacho-Sanchez, M., Velo-Anton, G., Hanson, J.O., Verissimo, A., Martinez-Solano, I., Marques, A., et al. (2020). Comparative assessment of range-wide patterns of genetic diversity and structure with SNPs and microsatellites: A case study with Iberian amphibians. Ecol Evol 10(19), 10353–10363. doi: 10.1002/ece3.6670.

Campino, S., Auburn, S., Kivinen, K., Zongo, I., Ouedraogo, J.B., Mangano, V., et al. (2011). Population genetic analysis of Plasmodium falciparum parasites using a customized Illumina GoldenGate genotyping assay. PLoS One 6(6), e20251. doi: 10.1371/journal.pone.0020251.

Dalmat, R., Naughton, B., Kwan-Gett, T.S., Slyker, J., and Stuckey, E.M. (2019). Use cases for genetic epidemiology in malaria elimination. Malar J 18(1), 163. doi: 10.1186/s12936-019-2784-0.

Daniels, R., Volkman, S.K., Milner, D.A., Mahesh, N., Neafsey, D.E., Park, D.J., et al. (2008). A general SNP-based molecular barcode for Plasmodium falciparum identification and tracking. Malar J 7, 223. doi: 10.1186/1475-2875-7-223.

Delgado-Ratto, C., Gamboa, D., Soto-Calle, V.E., Van den Eede, P., Torres, E., Sanchez-Martinez, L., et al. (2016). Population Genetics of Plasmodium vivax in the Peruvian Amazon. PLoS Negl Trop Dis 10(1), e0004376. doi: 10.1371/journal.pntd.0004376.

Escalante, A.A., Ferreira, M.U., Vinetz, J.M., Volkman, S.K., Cui, L., Gamboa, D., et al. (2015). Malaria Molecular Epidemiology: Lessons from the International Centers of Excellence for Malaria Research Network. Am J Trop Med Hyg 93(3 Suppl), 79–86. doi: 10.4269/ajtmh.15-0005.

Escalante, A.A., and Pacheco, M.A. (2019). Malaria Molecular Epidemiology: An Evolutionary Genetics Perspective. Microbiol Spectr 7(4). doi: 10.1128/microbiolspec.AME-0010-2019.

Evanno, G., Regnaut, S., and Goudet, J. (2005). Detecting the number of clusters of individuals using the software STRUCTURE: a simulation study. Mol Ecol 14(8), 2611–2620. doi: 10.1111/j.1365-294X.2005.02553.x.

Ferreira, M.U., Karunaweera, N.D., da Silva-Nunes, M., da Silva, N.S., Wirth, D.F., and Hartl, D.L. (2007). Population structure and transmission dynamics of Plasmodium vivax in rural Amazonia. J Infect Dis 195(8), 1218–1226. doi: 10.1086/512685.

Fola, A.A., Harrison, G.L.A., Hazairin, M.H., Barnadas, C., Hetzel, M.W., Iga, J., et al. (2017). Higher Complexity of Infection and Genetic Diversity of Plasmodium vivax Than Plasmodium falciparum Across All Malaria Transmission Zones of Papua New Guinea. Am J Trop Med Hyg 96(3), 630–641. doi: 10.4269/ajtmh.16-0716.

Fola, A.A., Kattenberg, E., Razook, Z., Lautu-Gumal, D., Lee, S., Mehra, S., et al. (2020). SNP barcodes provide higher resolution than microsatellite markers to measure Plasmodium vivax population genetics. Malar J 19(1), 375. doi: 10.1186/s12936-020-03440-0.

Francis, R.M. (2017). pophelper: an R package and web app to analyse and visualize population structure. Mol Ecol Resour 17(1), 27–32. doi: 10.1111/1755-0998.12509.

Ghansah, A., Tiedje, K.E., Argyropoulos, D.C., Onwona, C.O., Deed, S.L., Labbe, F., et al. (2023). Comparison of molecular surveillance methods to assess changes in the population genetics of Plasmodium falciparum in high transmission. Front Parasitol 2. doi: 10.3389/fpara.2023.1067966.

Golumbeanu, M., Edi, C.A.V., Hetzel, M.W., Koepfli, C., and Nsanzabana, C. (2023). Bridging the Gap from Molecular Surveillance to Programmatic Decisions for Malaria Control and Elimination. Am J Trop Med Hyg. doi: 10.4269/ajtmh.22-0749.

Goudet, J., Jombart, T., Kamvar, Z., Archer, E., and Hardy, O. (2022). Package ‘hierfstat’ [Online]. Available: https://cran.r-project.org/web/packages/hierfstat/hierfstat.pdf [Accessed].

Gwarinda, H.B., Tessema, S.K., Raman, J., Greenhouse, B., and Birkholtz, L.M. (2021). Parasite genetic diversity reflects continued residual malaria transmission in Vhembe District, a hotspot in the Limpopo Province of South Africa. Malar J 20(1), 96. doi: 10.1186/s12936-021-03635-z.

Harrison, G.L.A., Mehra, S., Razook, Z., Tessier, N., Lee, S., Hetzel, M.W., et al. (2024). Plasmodium falciparum populations, transmission dynamics and infection origins across Papua New Guinea. medRxiv, 2023.2009.2004.23294444. doi: 10.1101/2023.09.04.23294444.

Imwong, M., Nair, S., Pukrittayakamee, S., Sudimack, D., Williams, J.T., Mayxay, M., et al. (2007). Contrasting genetic structure in Plasmodium vivax populations from Asia and South America. Int J Parasitol 37(8-9), 1013–1022. doi: 10.1016/j.ijpara.2007.02.010.

Jolliffe, I.T., and Cadima, J. (2016). Principal component analysis: a review and recent developments. Philos Trans A Math Phys Eng Sci 374(2065), 20150202. doi: 10.1098/rsta.2015.0202.

Kanai, M., Yeo, T., Asua, V., Rosenthal, P.J., Fidock, D.A., and Mok, S. (2022). Comparative Analysis of Plasmodium falciparum Genotyping via SNP Detection, Microsatellite Profiling, and Whole-Genome Sequencing. Antimicrob Agents Chemother 66(1), e0116321. doi: 10.1128/AAC.01163-21.

Kattenberg, J.H., Cabrera-Sosa, L., Figueroa-Ildefonso, E., Mutsaers, M., Monsieurs, P., Guetens, P., et al. (2024). Plasmodium vivax genomic surveillance in the Peruvian Amazon with Pv AmpliSeq assay. PLoS Negl Trop Dis 18(7), e0011879. doi: 10.1371/journal.pntd.0011879.

Kattenberg, J.H., Fernandez-Minope, C., van Dijk, N.J., Llacsahuanga Allcca, L., Guetens, P., Valdivia, H.O., et al. (2023a). Malaria Molecular Surveillance in the Peruvian Amazon with a Novel Highly Multiplexed Plasmodium falciparum AmpliSeq Assay. Microbiol Spectr 11(2), e0096022. doi: 10.1128/spectrum.00960-22.

Kattenberg, J.H., Nguyen, H.V., Nguyen, H.L., Sauve, E., Nguyen, N.T.H., Chopo-Pizarro, A., et al. (2022). Novel highly-multiplexed AmpliSeq targeted assay for Plasmodium vivax genetic surveillance use cases at multiple geographical scales. Front Cell Infect Microbiol 12, 953187. doi: 10.3389/fcimb.2022.953187.

Kattenberg, J.H., Razook, Z., Keo, R., Koepfli, C., Jennison, C., Lautu-Gumal, D., et al. (2020). Monitoring Plasmodium falciparum and Plasmodium vivax using microsatellite markers indicates limited changes in population structure after substantial transmission decline in Papua New Guinea. Mol Ecol 29(23), 4525–4541. doi: 10.1111/mec.15654.

Kattenberg, J.H., Van Dijk, N.J., Fernandez-Minope, C.A., Guetens, P., Mutsaers, M., Gamboa, D., et al. (2023b). Molecular Surveillance of Malaria Using the PF AmpliSeq Custom Assay for Plasmodium falciparum Parasites from Dried Blood Spot DNA Isolates from Peru. Bio Protoc 13(5), e4621. doi: 10.21769/BioProtoc.4621.

Keenan, K., McGinnity, P., Cross, T.F., Crozier, W.W., and Prodöhl, P.A. (2013). diveRsity: An R package for the estimation and exploration of population genetics parameters and their associated errors. Methods in Ecology and Evolution 4(8), 782–788. doi: 10.1111/2041-210x.12067.

Kitada, S., Nakamichi, R., and Kishino, H. (2021). Understanding population structure in an evolutionary context: population-specific FST and pairwise FST. G3 (Bethesda) 11(11). doi: 10.1093/g3journal/jkab316.

Lo, E., Bonizzoni, M., Hemming-Schroeder, E., Ford, A., Janies, D.A., James, A.A., et al. (2018). Selection and Utility of Single Nucleotide Polymorphism Markers to Reveal Fine-Scale Population Structure in Human Malaria Parasite. Frontiers in Ecology and Evolution 6. doi: 10.3389/fevo.2018.00145.

Lopez, L., and Koepfli, C. (2021). Systematic review of Plasmodium falciparum and Plasmodium vivax polyclonal infections: Impact of prevalence, study population characteristics, and laboratory procedures. PLoS One 16(6), e0249382. doi: 10.1371/journal.pone.0249382.

Mangold, K.A., Manson, R.U., Koay, E.S., Stephens, L., Regner, M., Thomson, R.B., Jr., et al. (2005). Real-time PCR for detection and identification of Plasmodium spp. J Clin Microbiol 43(5), 2435–2440. doi: 10.1128/JCM.43.5.2435-2440.2005.

Manrique, P., Hoshi, M., Fasabi, M., Nolasco, O., Yori, P., Calderon, M., et al. (2015). Assessment of an automated capillary system for Plasmodium vivax microsatellite genotyping. Malar J 14, 326. doi: 10.1186/s12936-015-0842-9.

Manrique, P., Miranda-Alban, J., Alarcon-Baldeon, J., Ramirez, R., Carrasco-Escobar, G., Herrera, H., et al. (2019). Microsatellite analysis reveals connectivity among geographically distant transmission zones of Plasmodium vivax in the Peruvian Amazon: A critical barrier to regional malaria elimination. PLoS Negl Trop Dis 13(11), e0007876. doi: 10.1371/journal.pntd.0007876.

Mathema, V.B., Nakeesathit, S., White, N.J., Dondorp, A.M., and Imwong, M. (2020). Genome-wide microsatellite characteristics of five human Plasmodium species, focusing on Plasmodium malariae and P. ovale curtisi. Parasite 27, 34. doi: 10.1051/parasite/2020034.

Meirmans, P.G., and Hedrick, P.W. (2011). Assessing population structure: F(ST) and related measures. Mol Ecol Resour 11(1), 5–18. doi: 10.1111/j.1755-0998.2010.02927.x.

Ministerio de Salud (2015). Norma técnica de salud para la atención de la malaria y malaria grave en el Perú [Online]. Available: http://bvs.minsa.gob.pe/local/MINSA/4373.pdf [Accessed].

Murray, L., Mobegi, V.A., Duffy, C.W., Assefa, S.A., Kwiatkowski, D.P., Laman, E., et al. (2016). Microsatellite genotyping and genome-wide single nucleotide polymorphism-based indices of Plasmodium falciparum diversity within clinical infections. Malar J 15(1), 275. doi: 10.1186/s12936-016-1324-4.

Nair, S., Williams, J.T., Brockman, A., Paiphun, L., Mayxay, M., Newton, P.N., et al. (2003). A selective sweep driven by pyrimethamine treatment in southeast asian malaria parasites. Mol Biol Evol 20(9), 1526–1536. doi: 10.1093/molbev/msg162.

Nderu, D., Kimani, F., Karanja, E., Thiong’o, K., Akinyi, M., Too, E., et al. (2019). Genetic diversity and population structure of Plasmodium falciparum in Kenyan-Ugandan border areas. Trop Med Int Health 24(5), 647–656. doi: 10.1111/tmi.13223.

Neafsey, D.E., Taylor, A.R., and MacInnis, B.L. (2021). Advances and opportunities in malaria population genomics. Nat Rev Genet 22(8), 502–517. doi: 10.1038/s41576-021-00349-5.

Nei, M., and Roychoudhury, A.K. (1974). Sampling variances of heterozygosity and genetic distance. Genetics 76(2), 379–390. doi: 10.1093/genetics/76.2.379.

Noviyanti, R., Miotto, O., Barry, A., Marfurt, J., Siegel, S., Thuy-Nhien, N., et al. (2020). Implementing parasite genotyping into national surveillance frameworks: feedback from control programmes and researchers in the Asia-Pacific region. Malar J 19(1), 271. doi: 10.1186/s12936-020-03330-5.

Padilla-Rojas, C., Jimenez-Vasquez, V., Hurtado, V., Mestanza, O., Molina, I.S., Barcena, L., et al. (2021). Genomic analysis reveals a rapid spread and predominance of lambda (C.37) SARS-COV-2 lineage in Peru despite circulation of variants of concern. J Med Virol 93(12), 6845–6849. doi: 10.1002/jmv.27261.

Pava, Z., Puspitasari, A.M., Rumaseb, A., Handayuni, I., Trianty, L., Utami, R.A.S., et al. (2020). Molecular surveillance over 14 years confirms reduction of Plasmodium vivax and falciparum transmission after implementation of Artemisinin-based combination therapy in Papua, Indonesia. PLoS Negl Trop Dis 14(5), e0008295. doi: 10.1371/journal.pntd.0008295.

Peakall, R., and Smouse, P.E. (2012). GenAlEx 6.5: genetic analysis in Excel. Population genetic software for teaching and research--an update. Bioinformatics 28(19), 2537–2539. doi: 10.1093/bioinformatics/bts460.

Perez-Gonzalez, J., Carranza, J., Anaya, G., Broggini, C., Vedel, G., de la Pena, E., et al. (2023). Comparative Analysis of Microsatellite and SNP Markers for Genetic Management of Red Deer. Animals (Basel) 13(21). doi: 10.3390/ani13213374.

Pritchard, J.K., Stephens, M., and Donnelly, P. (2000). Inference of population structure using multilocus genotype data. Genetics 155(2), 945–959. doi: 10.1093/genetics/155.2.945.

Puckett, E.E., and Eggert, L.S. (2016). Comparison of SNP and microsatellite genotyping panels for spatial assignment of individuals to natal range: A case study using the American black bear (*Ursus americanus*). Biological Conservation 193, 86–93. doi: 10.1016/j.biocon.2015.11.020.

Puyen, Z.M., Santos-Lazaro, D., Vigo, A.N., Cotrina, V.V., Ruiz-Nizama, N., Alarcon, M.J., et al. (2024). Whole Genome Sequencing of Mycobacterium tuberculosis under routine conditions in a high-burden area of multidrug-resistant tuberculosis in Peru. PLoS One 19(6), e0304130. doi: 10.1371/journal.pone.0304130.

Roper, C., Pearce, R., Bredenkamp, B., Gumede, J., Drakeley, C., Mosha, F., et al. (2003). Antifolate antimalarial resistance in southeast Africa: a population-based analysis. Lancet 361(9364), 1174–1181. doi: 10.1016/S0140-6736(03)12951-0.

Rosado, J., Carrasco-Escobar, G., Nolasco, O., Garro, K., Rodriguez-Ferruci, H., Guzman-Guzman, M., et al. (2022). Malaria transmission structure in the Peruvian Amazon through antibody signatures to Plasmodium vivax. PLoS Negl Trop Dis 16(5), e0010415. doi: 10.1371/journal.pntd.0010415.

Rougemont, M., Van Saanen, M., Sahli, R., Hinrikson, H.P., Bille, J., and Jaton, K. (2004). Detection of four Plasmodium species in blood from humans by 18S rRNA gene subunit-based and species-specific real-time PCR assays. J Clin Microbiol 42(12), 5636–5643. doi: 10.1128/JCM.42.12.5636-5643.2004.

Rovira-Vallbona, E., Kattenberg, J.H., Hong, N.V., Guetens, P., Imamura, H., Monsieurs, P., et al. (2023a). Molecular surveillance of Plasmodium falciparum drug-resistance markers in Vietnam using multiplex amplicon sequencing (2000-2016). Sci Rep 13(1), 13948. doi: 10.1038/s41598-023-40935-7.

Rovira-Vallbona, E., Kattenberg, J.H., Van Hong, N., Guetens, P., Imamura, H., Monsieurs, P., et al. (2023b). Author Correction: Molecular surveillance of Plasmodium falciparum drug-resistance markers in Vietnam using multiplex amplicon sequencing (2000-2016). Sci Rep 13(1), 17207. doi: 10.1038/s41598-023-43996-w.

Ruybal-Pesantez, S., McCann, K., Vibin, J., Siegel, S., Auburn, S., and Barry, A.E. (2024). Molecular markers for malaria genetic epidemiology: progress and pitfalls. Trends Parasitol 40(2), 147–163. doi: 10.1016/j.pt.2023.11.006.

Salado, I., Fernandez-Gil, A., Vila, C., and Leonard, J.A. (2021). Automated genotyping of microsatellite loci from feces with high throughput sequences. PLoS One 16(10), e0258906. doi: 10.1371/journal.pone.0258906.

Schaffner, S.F., Badiane, A., Khorgade, A., Ndiop, M., Gomis, J., Wong, W., et al. (2023). Malaria surveillance reveals parasite relatedness, signatures of selection, and correlates of transmission across Senegal. Nat Commun 14(1), 7268. doi: 10.1038/s41467-023-43087-4.

Serrote, C.M.L., Reiniger, L.R.S., Silva, K.B., Rabaiolli, S., and Stefanel, C.M. (2020). Determining the Polymorphism Information Content of a molecular marker. Gene 726, 144175. doi: 10.1016/j.gene.2019.144175.

Sutton, P.L. (2013). A call to arms: on refining Plasmodium vivax microsatellite marker panels for comparing global diversity. Malar J 12, 447. doi: 10.1186/1475-2875-12-447.

Sy, M., Deme, A.B., Warren, J.L., Early, A., Schaffner, S., Daniels, R.F., et al. (2022). Plasmodium falciparum genomic surveillance reveals spatial and temporal trends, association of genetic and physical distance, and household clustering. Sci Rep 12(1), 938. doi: 10.1038/s41598-021-04572-2.

Taylor, A.R., Jacob, P.E., Neafsey, D.E., and Buckee, C.O. (2019). Estimating Relatedness Between Malaria Parasites. Genetics 212(4), 1337–1351. doi: 10.1534/genetics.119.302120.

Tessema, S., Wesolowski, A., Chen, A., Murphy, M., Wilheim, J., Mupiri, A.R., et al. (2019). Using parasite genetic and human mobility data to infer local and cross-border malaria connectivity in Southern Africa. Elife 8. doi: 10.7554/eLife.43510.

Theissinger, K., Fernandes, C., Formenti, G., Bista, I., Berg, P.R., Bleidorn, C., et al. (2023). How genomics can help biodiversity conservation. Trends Genet 39(7), 545–559. doi: 10.1016/j.tig.2023.01.005.

Trimarsanto, H., Amato, R., Pearson, R.D., Sutanto, E., Noviyanti, R., Trianty, L., et al. (2022). A molecular barcode and web-based data analysis tool to identify imported Plasmodium vivax malaria. Commun Biol 5(1), 1411. doi: 10.1038/s42003-022-04352-2.

Villasis, E., Garro, K., Rosas-Aguirre, A., Rodriguez, P., Rosado, J., Gave, A., et al. (2021). PvMSP8 as a Novel Plasmodium vivax Malaria Sero-Marker for the Peruvian Amazon. Pathogens 10(3). doi: 10.3390/pathogens10030282.

Wamae, K., Kimenyi, K.M., Osoti, V., de Laurent, Z.R., Ndwiga, L., Kharabora, O., et al. (2022). Amplicon Sequencing as a Potential Surveillance Tool for Complexity of Infection and Drug Resistance Markers in Plasmodium falciparum Asymptomatic Infections. J Infect Dis 226(5), 920–927. doi: 10.1093/infdis/jiac144.

Wasakul, V., Disratthakit, A., Mayxay, M., Chindavongsa, K., Sengsavath, V., Thuy-Nhien, N., et al. (2023). Malaria outbreak in Laos driven by a selective sweep for Plasmodium falciparum kelch13 R539T mutants: a genetic epidemiology analysis. Lancet Infect Dis 23(5), 568–577. doi: 10.1016/S1473-3099(22)00697-1.

Weir, B.S., and Cockerham, C.C. (1984). Estimating F-Statistics for the Analysis of Population Structure. Evolution 38(6), 1358–1370. doi: 10.1111/j.1558-5646.1984.tb05657.x.

World Health Organization (2021). Global technical strategy for malaria 2016-2030, 2021 update [Online]. Geneva: World Health Organization. Available: https://www.who.int/publications/i/item/9789240031357 [Accessed].

World Health Organization (2023). World malaria report 2023 [Online]. Geneva: World Health Organization. Available: https://www.who.int/teams/global-malaria-programme/reports/world-malaria-report-2023 [Accessed].

Zhong, D., Koepfli, C., Cui, L., and Yan, G. (2018). Molecular approaches to determine the multiplicity of Plasmodium infections. Malar J 17(1), 172. doi: 10.1186/s12936-018-2322-5.

Zimmerman, S.J., Aldridge, C.L., and Oyler-McCance, S.J. (2020). An empirical comparison of population genetic analyses using microsatellite and SNP data for a species of conservation concern. BMC Genomics 21(1), 382. doi: 10.1186/s12864-020-06783-9.

