## Supplementary material for "Comparing newly developed SNP barcode panels with microsatellites to explore population genetics of malaria parasites in the Peruvian Amazon": Suplementary Material

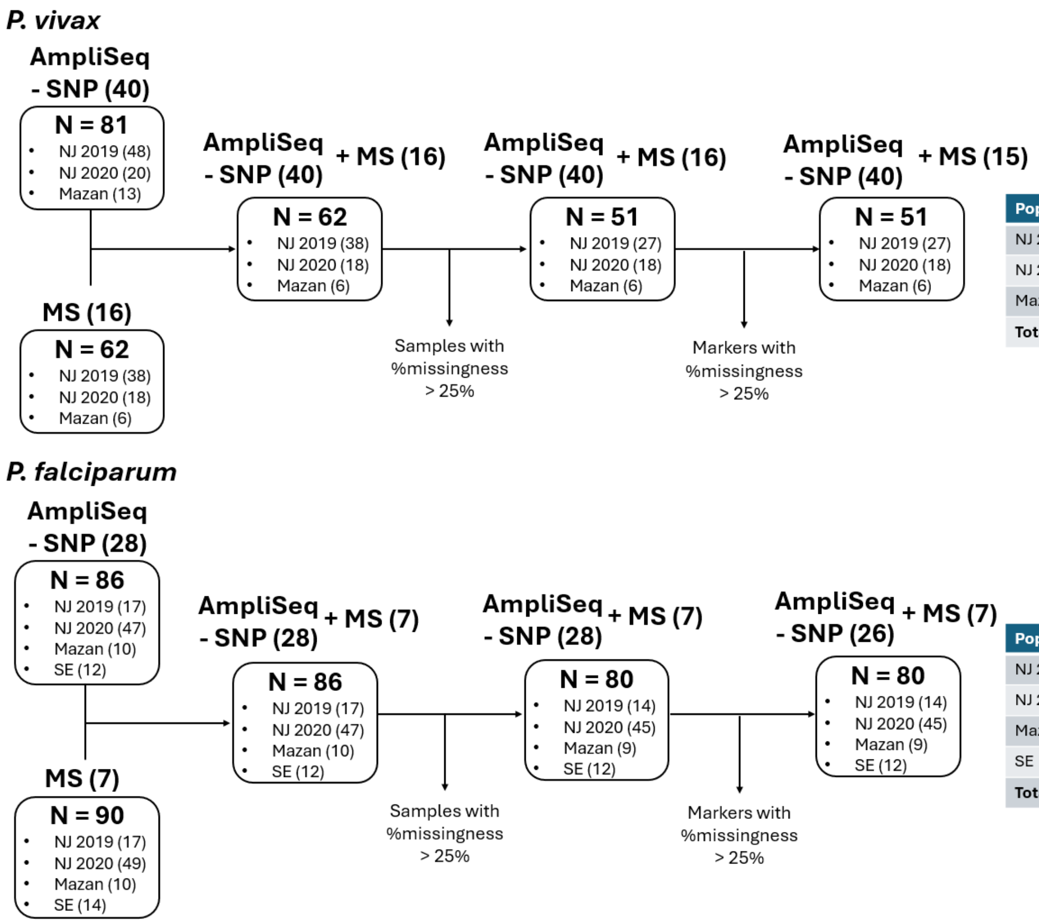


| Pop | MLG - SNP | MLG - MS |
|---|---|---|
| NJ 2019 | 35 | 45 |
| NJ 2020 | 25 | 32 |
| Mazan | 8 | 9 |
| **Total** | **68** | **86** |

| Pop | MLG - SNP | MLG - MS |
|---|---|---|
| NJ 2019 | 22 | 18 |
| NJ 2020 | 72 | 65 |
| Mazan | 9 | 10 |
| SE | 14 | 12 |
| **Total** | **117** | **105** |

**Supplementary Figure S1. Flowchart describing number and source of Pv and Pf samples and inclusion criteria.** The tables at the right show the number of haplotypes included in the analyzes. MS: microsatellite, NJ: Nueva Jerusalen, SE: Santa Emilia, Pop: population, MLG: N° of multi-locus genotypes.

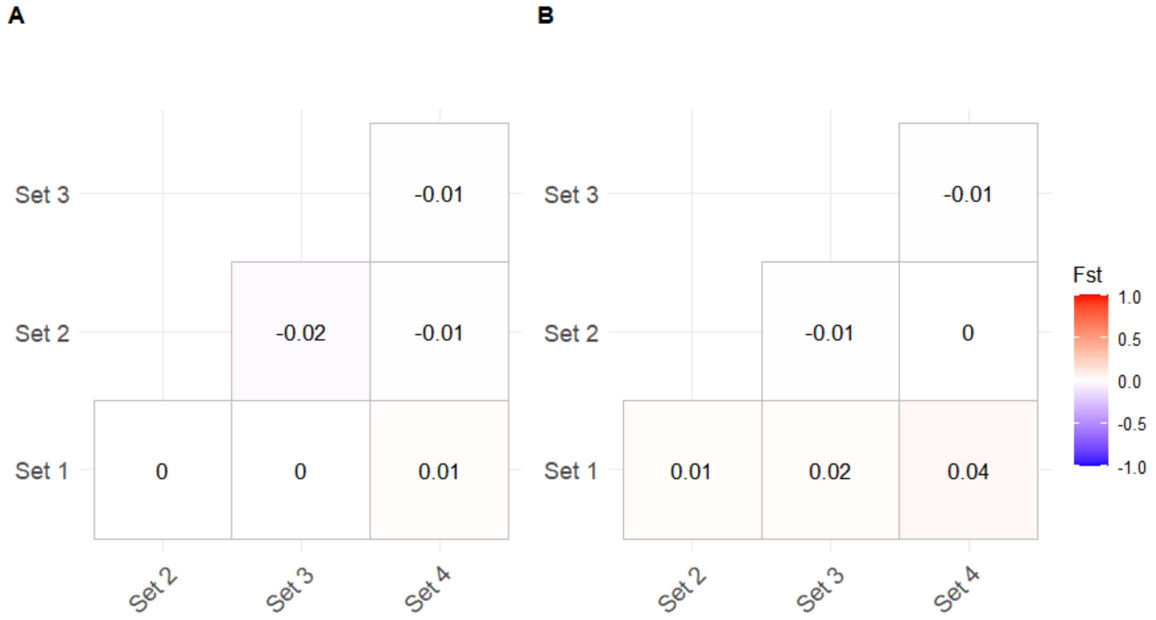


**Supplementary Figure S2. Inclusion of secondary MLG.** Heatmap of pairwise FST values between sets containing different type of MLG. Set 1: samples with only one MLG. Set 2: dominant MLG of all samples. Set 3: dominant MLG of all samples and secondary MLG of samples with in only one secondary allele. Set 4: all dominant and secondary MLG of all samples

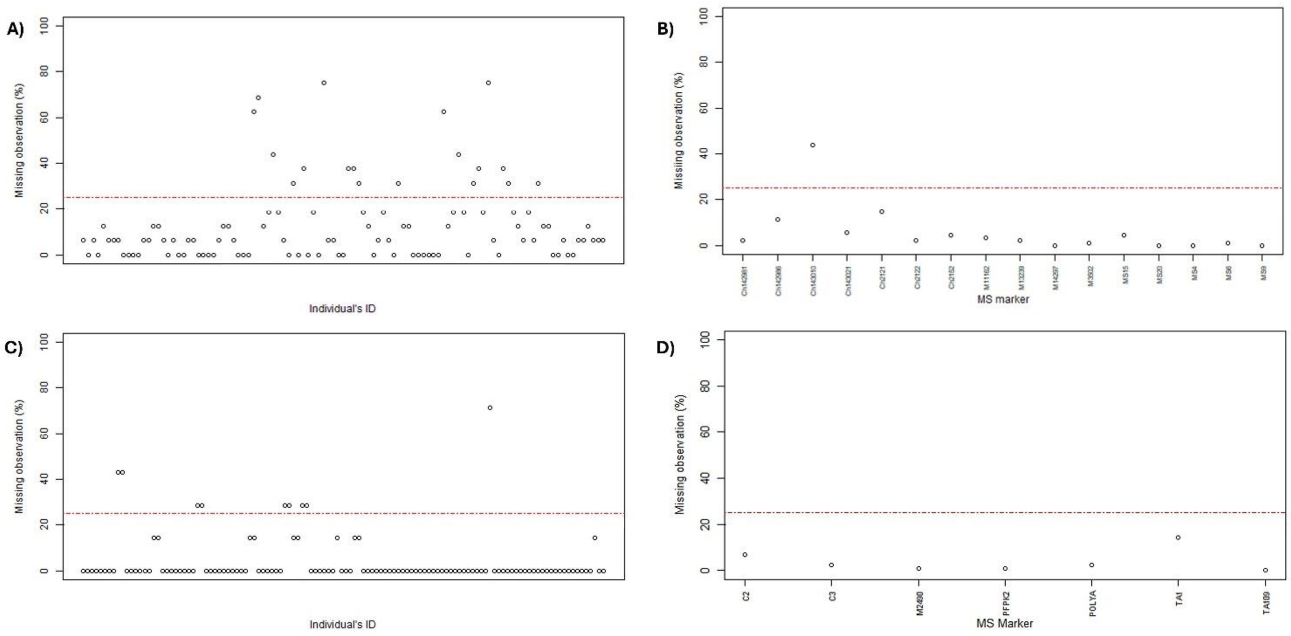


**Supplementary Figure S3. MLG and MS markers excluded from the analysis.** The red dashed line indicates missing rate threshold of 25%. The x-axis corresponds to the individual MLG identification (ID) (A and C) or the names of MS markers (B and D). (A) MLG from MS genotyping in Pv (A) and Pf (B) populations. MS panels for Pv (B) and Pf (D).

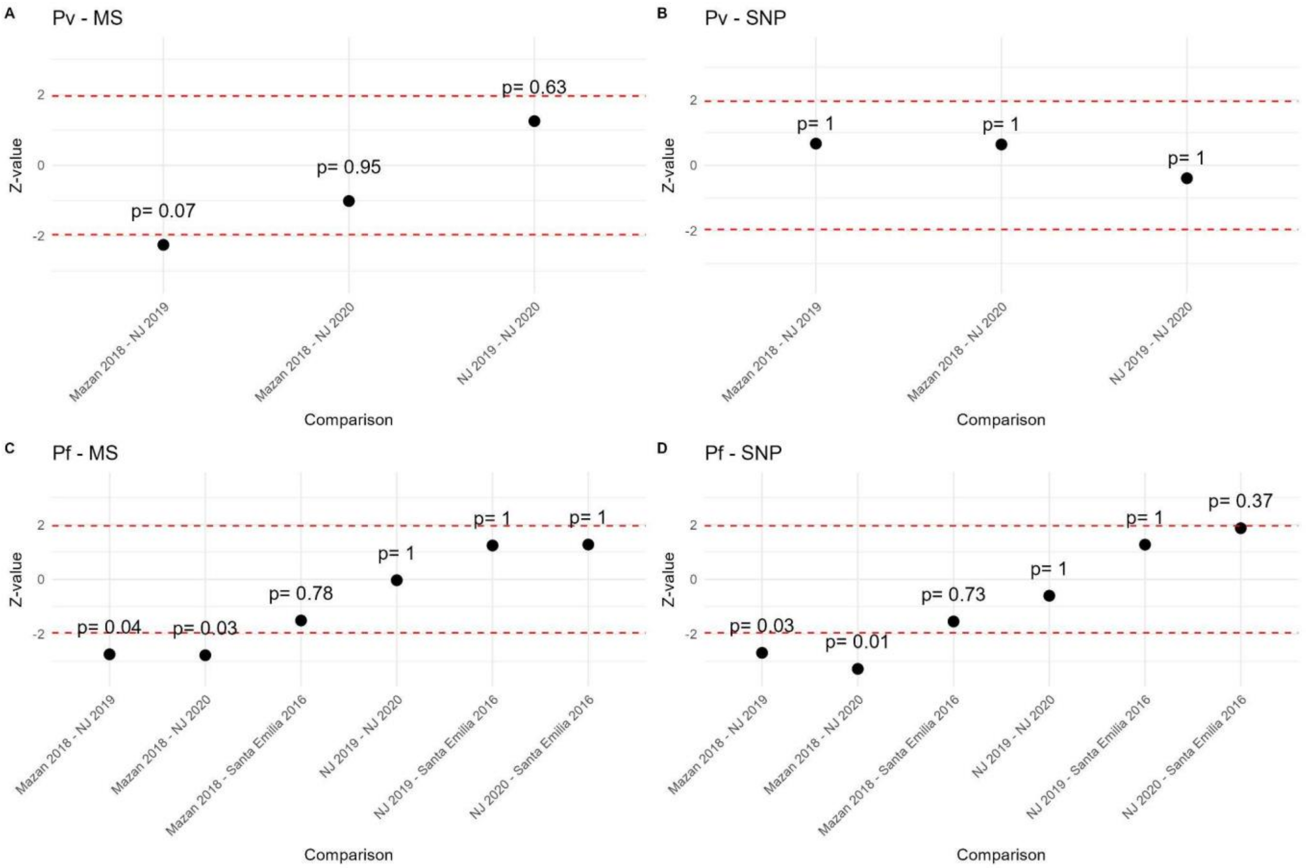


**Supplementary Figure S4. Dunn's post hoc comparisons of genetic diversity (expressed as expected heterozygosity, He) across Pv and Pf populations**. The red dashed lines represent z-values of -1.96 and 1.96, corresponding to a 5% significance level. The black circles indicate the z-scores calculated for each paired comparison, with the associated adjusted p-values displayed alongside each dot. The post hoc tests were performed using He values obtained with (A) MS panel in Pv, (B) SNP barcode for Pv, (C) MS panel in Pf, (D) SNP barcode for Pf. No significant differences were found in Pv with any panel. In Pf, only the comparisons between Mazan and NJ (2019 & 2020) were significant across both panels.

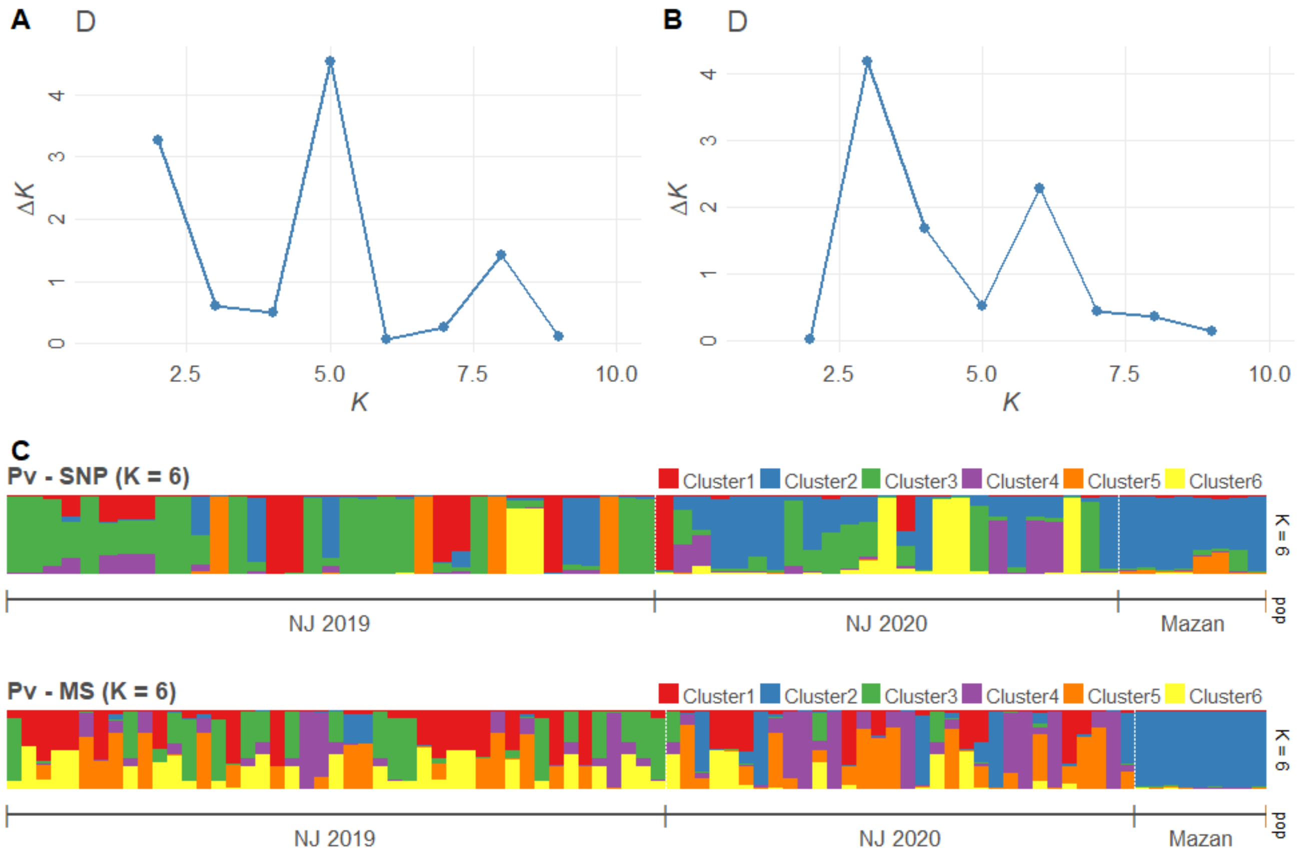


**Supplementary Figure S5.** Evanno plots of the STRUCTURE analysis in Pv samples using **(A)** SNP or **(B)** MS. **(C)** Clustering analysis by STRUCTURE at K = 7 using SNP or MS in Pv populations by location.

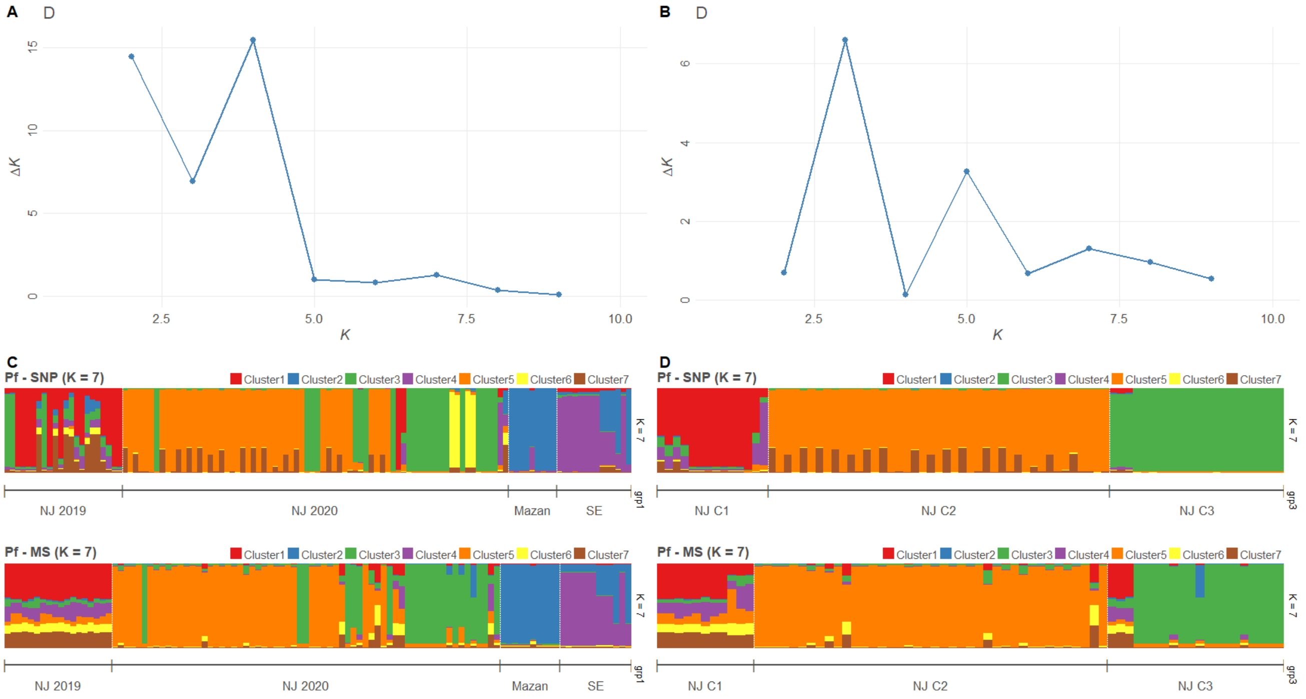


**Supplementary Figure S6.** Evanno plots of the STRUCTURE analysis in Pf samples using **(A)** SNP or **(B)** MS. Clustering analysis by STRUCTURE at K = 7 using SNP or MS in **(C)** Pf populations by location or **(D)** Pf samples in NJ by NJ by previous classification according to Cabrera-Sosa et al. (2024).

**Supplementary Table**

**Supplementary Table S1. Pv and Pf sample size and collection data used in this study from different previous projects.**

| Project (SIDISI code) | Area | Year of collection | Type of collection | *P. vivax* | | | *P. falciparum* | | |
|---|---|---|---|---|---|---|---|---|---|
| | | | | **Sample size - AmpliSeq** | **Sample size - MS** | **Final Sample size** | **Sample size - AmpliSeq** | **Sample size - MS** | **Final Sample size** |
| VLIR-TEAM (102725) | NJ | 2019 | ACD | 48 | 38 | **38** | 17 | 17 | **17** |
| | | 2020 | PCD | 20 | 18 | **16** | 47 | 49 | **47** |
| ICERM 2.0 (101518) | Mazan | 2018 | Population-based survey | 13 | 6 | **6** | 10 | 10 | **10** |
| | SE | 2016 | ACD & PCD | - | - | - | 12 | 14 | **12** |

NJ: Nueva Jerusalen, SE: Santa Emilia, ACD: active case detection, PCD: passive case detection

**Supplementary Table S2. Fixed positions in the SNP barcodes for Pv and Pf populations.**

| **Fixed positions in Pv Peru SNP barcode** |
| --- |
| PvP01_03_v1_114702 |
| PvP01_07_v1_959564 |
| **Fixed positions in Pf Peru SNP barcode** |
| Pf3D7_03_v3_849476 |
| Pf3D7_05_v3_921893 |
| Pf3D7_06_v3_636044 |
| Pf3D7_07_v3_782111 |
| Pf3D7_08_v3_803172 |
| Pf3D7_10_v3_1172712 |
| Pf3D7_10_v3_341106 |
| Pf3D7_11_v3_1505533 |
| Pf3D7_12_v3_1552084 |
| Pf3D7_14_v3_1381943 |
